# DNA shape and epigenomics distinguish the mechanistic origin of human genomic structural variations

**DOI:** 10.1101/2025.09.20.677549

**Authors:** Nadejda B. Boev, Mark B. Gerstein, Sushant Kumar

## Abstract

The recent advent of long-read whole genome sequencing has enabled us to create an accurate telomere-to-telomere reference genome, construct pangenome graphs, and compile precise catalogs of genomic structural variations (SVs). These comprehensive SV repositories provide an excellent opportunity to explore the role of SVs in genotype-phenotype associations and examine the mechanisms by which SVs are introduced through double-strand break (DSB) repair. Here, we employed comprehensive SV catalogs identified through various short- and long-read whole genome sequencing efforts to infer the underlying mechanisms of SV introduction based on their genomic and epigenomic profiles. Our findings indicate that high local DNA methylation and DNA shape-related features, such as low variations in propeller twist, support the origins of homology-driven SVs. Subsequently, we utilized an active-learning-based unsupervised clustering approach, revealing that the homology-dependent SVs show greater evidence of retaining ancestral recombination patterns compared to their homology-independent counterparts. Finally, our comparison of inherited and de novo SVs from healthy populations and rare disease cohorts showed distinct upstream H3K27me3 levels in de novo SVs from individuals with ultra-rare disorders. These findings highlight genome-wide characteristics that may influence the choice of repair mechanisms linked to heritable SV origins.

## Introduction

Over the last decade, there has been a continuous increase in large-scale genomic sequencing studies aimed at investigating genotype-phenotype relationships across healthy populations and various common and rare diseases(1, 2). Most of these studies have focused on characterizing short genomic variants, including single-nucleotide variants (SNVs) and small insertions/deletions (INDELs) under 50 bp. However, structural variants (SVs), operationally defined as variants longer than 50 bp, represent a significant source of genomic variation, as they induce more nucleotide-level changes than both SNVs and INDELs combine(3). SVs are further categorized as insertions, deletions, inversions, and translocations based on their orientation and modifications to DNA segments relative to a reference genome. Due to their longer length, SVs can simultaneously influence multiple coding and regulatory elements(4, 5) and are implicated in various diseases(6–8).

Current genomic studies primarily focus on interpreting SNVs and INDELs due to our superior ability to identify these variants accurately. Short-read whole genome sequencing (srWGS) platforms are commonly employed in large-scale genomic studies; however, they struggle to produce a precise SV catalog. This challenge stems from SVs occurring in or near highly repetitive regions of the human genome, resulting in multi-mapping and/or mis-mapping of sequencing reads, which leads to inaccurate or non-specific SV calling(9, 10). The recent emergence of long-read whole genome sequencing (lrWGS) aims to address these challenges. Implementing lrWGS in large-scale studies has generated comprehensive SV maps for healthy populations and various diseased individuals, deepening our understanding of population diversity(11–13) and uncovering the pathogenic role of SVs in diseases(7, 14). These new SV repositories also present a valuable opportunity to investigate the mechanistic origins of SVs.

Previous studies have linked the origin of SVs to double-strand breaks (DSBs) resulting from either effective or ineffective repair attempts(15–18). These mechanisms include non-homologous end joining (NHEJ), alternative end-joining (aEJ), single-strand annealing (SSA), and homologous recombination (HR), which require increasing amounts of local homology and resection during their repair process(15). Importantly, faulty repair mechanisms can lead to various diseases. For example, global DSB repair deficiencies are associated with LIG4 syndrome(19), Fanconi anemia(20), and an increased risk of multiple cancers(21). Additionally, defective non-allelic homologous recombination (NAHR) can result in 15q13.3 microdeletion syndrome(22). Finally, evaluating DSB repair efficacy can help predict therapeutic responses in patients with inherited or acquired HR deficiency (HRD) when receiving treatment with PARP inhibitors for breast, ovarian, prostate, or pancreatic cancers(23, 24).

Considering their clinical utility, computational and experimental studies have sought to understand DSBs and the mechanisms by which DSB repair can introduce variants (including SVs). For instance, previous research has employed repair-associated features to construct simple decision trees for categorizing germline SVs found in the 1000 Genomes Project (1KG phase 1)(25, 26) and variants in HuRef(27). While this approach is sound, these methods mainly rely on sequence homology and repetitive element annotations to define an SV mechanism class. Moreover, these studies have only characterized a small subset of SVs derived from low-coverage srWGS data, thereby overlooking the comprehensive landscape of SVs across the genome. In contrast, in recent years, numerous experimental studies have revealed complex interactions among various genomic, epigenomic, and biophysical features during DSB repair(28, 29). For example, multiple low-throughput reporter assays (such as BLESS(30) or TRIP(31)) suggest that DNA accessibility is a relevant repair feature(16, 32, 33). However, these studies are inherently biased(34), as DSB formation is influenced by the selected restriction enzyme or CRISPR(35, 36), variability in mechanism frequency due to the lack of cell cycle synchronization(37), and biased repair usage depending on the cell types and states(38, 39). The inherent limitations of experimental approaches have resulted in inconsistencies across these studies. Furthermore, structural biology studies have indicated that the biophysical features of DNA near the breakpoints, such as flexibility, kinking, or minor groove width (MGW), may lead to preferential repair by repair complexes, although these studies are limited in throughput (40, 41). Thus, it remains unclear whether there are strict rules governing DSB repair choices. For example, whether euchromatin or heterochromatin, including facultative and constitutive, is consistently and faithfully repaired via HR or another homology-based repair mechanism (ie. SSA or aEJ)(42).

Considering these knowledge gaps and the rise of lrWGS-based accurate SV catalogs, sufficient data is now available to revisit and explore the relationship between SVs and features that support DSB repair on a genome-wide scale. Therefore, we utilized accurately identified SVs from multiple lrWGS efforts (including the Human Genome Structural Variation Consortium(13) (HGSVC) and 1KG-ONT(43) study, as well as high-coverage srWGS (1KG phase3 extended project)(44) to comprehensively characterize the features underlying repair mechanisms. We began similarly to previous work(13, 25, 26) by employing a straightforward homology-based workflow to categorize SVs into distinct mechanism classes. Then, we systematically investigated the underlying discriminatory sequence, biophysical, and epigenomic patterns associated with different homology-based SV categories identified by our simple workflow. For instance, we found that local DNA shape and certain genome accessibility features can differentiate SV classes. Subsequently, we trained active-learning-based unsupervised machine learning models to exploit these discriminative features, going beyond sequence homology to refine a given SV’s underlying repair mechanism class. Next, we applied these models to rare SVs from the high coverage 1KG project to recapitulate the established relationship between recombination rates and repair mechanisms in an ancestry-specific manner. Finally, a methodical comparison of *de novo* SVs from rare diseases and the 1KG dataset revealed differential levels of the H3K27me3 histone modification. Overall, our work develops and applies new methods to provide extensive insights into SVs and their mechanisms of introduction.

## Materials & Methods

We utilized resolved germline SVs from the HGSVC2, 1KG, and 1KG-ONT projects in this work. All these studies employed the hg38 reference genome to identify variants. We note that sequencing, assembly algorithms, and variant calling strategies were project-specific. HGSVC2 serves as a resource of haplotype-specific SVs identified with Phase Assembly Variant (PAV)(13) and merged across 35 individuals sequenced with PacBio (unique SVs: n_INS_= 65,836; n_DEL_=40,496). The pilot 1KG-ONT study identified SVs from Oxford Nanopore sequencing involving 100 individuals. The SVs were merged using Jasmine(83) and subject to five distinct SV calling strategies (unique SVs: n_INS_= 62,929, n_DEL_= 49,435). Finally, the 1KG study contains SVs from 3,202 individuals sequenced with Illumina, which includes 602 trios. We used the high-confidence variants generated by integrating SVs using GATK-SV(84), svtools(85), and Absinthe (https://github.com/nygenome/absinthe; unique SVs: n_DEL_=57,243). For HGSVC2 and 1KG-ONT, we utilized both insertions and deletions, while for the 1KG project, we only used deletions to mitigate quality issues from assembling insertions from short-read sequencing.

### Building a homology-based workflow

We initially labeled SVs solely based on their local homology. Using the reference genome, we extracted 1975 bp sequences upstream and downstream of the breakpoints, which were concatenated to the first or last 25bp of the SVs, respectively. Next, we performed a gapless alignment between these two sequences with a word size of 5 and a minimum identity of 80% using BLAST. We required alignments to have opposing orientations and to include ±2bp of the breakpoint. In this way, we considered alignments that captured plausible repair events via the introduction of the SV. Finally, we selected the alignment result with the highest bit score. Our goal was to create SV classes reflective reflect DSB repair pathways, including HR, SSA, aEJ, and NHE(46–49). Therefore, we classified SVs with long alignments (a minimum length of 100 bp out of 2000 bp) and high-quality alignments (a minimum identity of 90%) into the high local homology (HLH) class, inspired by HR. In contrast, we designated SVs to the intermediate local homology (ILH) class if they had shorter alignments (length between 20 and 100 bp) and slightly lower quality alignments (minimum identity of 80%). Next, we labeled variants for which BLAST could not identify homology as having no local homology (NLH), representative of SVs associated with NHEJ. Lastly, we categorized SVs where homology was recognized but did not meet the HLH or ILH thresholds as undefined. To describe the concordance between this workflow and Breakseq v2(25), we assigned HGSVC2 SVs using both methods.

To demonstrate the efficacy of this workflow, we utilized somatic SVs from cancer cohorts. We hypothesized that since extreme variant patterns are not anticipated within the germline variation profile of a healthy genome, utilizing samples with known deficiencies or proficiencies in relevant repair pathways would provide a better proxy for evaluating our framework. Therefore, we obtained SVs sequenced using Illumina and called with Manta(86) from the 100KG Project(50). We retrieved mutational signature contributions previously computed using COSMICv2(87) in tandem. We established cohorts to represent positive and negative controls based on evidence of HR deficiency (HRD) or proficiency (HRP) and mismatch repair deficiency (MMRD) or proficiency (MMRP), as their mechanisms differ in their utilization of homology. We calculated the proportion of deletions assigned to NLH within a sample. We considered the workflow successful if HRD and HRP samples exhibited a noticeable shift in %NLH, while MMRD and MMRP samples did not. We selected breast invasive carcinoma (BRCA) and colon adenocarcinoma (COAD) because of their higher rates of HHRD and MMRD, respectively(51), where we utilized signatures 3 and 6, respectively. We defined a sample as deficient in a repair mechanism if the relevant signature’s contribution was above the 95th percentile within the cancer study. Conversely, samples with contributions of 0 in the appropriate signature were considered “proficient." Consequently, we identified 152 samples with HRD, 909 with HRP, 103 with MMRD, and 155 with MMRP.

### Active-learning-based unsupervised clustering strategy

We aimed to develop a model that better capitalizes on the features underlying DSB repair, extending beyond just local homology. Therefore, we annotated SVs, their breakpoints, and their 2000 bp upstream and downstream flanking sequences with features including sequence characteristics, epigenomics, replication timing, repetitive elements, non-B DNA structures, and DNA shape. Given the chromosome and length biases across these features, we generated 100 simulations using SURVIVOR(54), which matched the real SVs in type, chromosome, and length. Subsequently, we annotated both the observed and simulated SV datasets.

The sequence features we calculated for the SVs and flanks included means and standard deviations in %GC content, stability, flexibility, and complexity (Shannon’s entropy with 10bp sliding windows). When annotating germline SVs, we used data from H1-hESC as a proxy due to its spatial and temporal similarities in introducing SVs in the germline(53). We estimated epigenomic profiles via Chip-Seq (CTCF, H2AFZ, H3K27ac, H3K27me3, H3K36me3, H3K4me1, H3K79me2, H3K9ac, H3K9me3, H4K20me1), DNase-seq, and WGB-seq experiments provided by ENCODE. We downloaded datasets from the ENCODE portal(88) (https://www.encodeproject.org/) with the following identifiers: ENCBS718AAA and ENCBS111ENC. We calculated the mean and standard deviation of the signal for the flanks (insertions and deletions) and the deleted regions using pybigwig (https://github.com/deeptools/pyBigWig/). We used a 16-stage Repli-seq experiment(89) from the 4D Nucleome Data portal(90) to quantify the replication timing at SV breakpoints. This data was further processed using the RepliSeq package(91), which provides an S50 value. S50 represents the S phase fraction, between 0 (early) and 1 (late), indicating the fraction of cells in a region that have undergone replication. To describe repetitive elements, we counted the number of LINEs, low complexity regions, LTRs, satellites, simple repeats, and SINE motifs in the flanks and SV, as determined by RepeatMasker. Similarly, we counted the number of non-B DNA structures (G-quadruplexes, Z-DNA, A-phased repeats, direct repeats, mirrored repeats, and short tandem repeats) within the flanking sequences using the nonB-GFA tool(92). Finally, we calculated the mean and standard deviation of DNA shape (minor groove width (MGW), helical twist (HelT), propeller twist (ProT), roll, and electrostatic potential (EP)) within the SV and flanking sequences using DNAShapeR(93). Briefly, the DNA shape features produced by DNAShapeR were derived from pentamer nucleotides that underwent all-atom Monte Carlo (MC) simulations. This pentamer dataset was compiled by conducting MC simulations of 2121 distinct DNA fragments of varying lengths (12-27 bp) sourced from DNA and protein-DNA complex structures available in the Protein Data Bank. Upon completing the annotations, we matched the real SVs to their simulations to calculate a z-score normalized value for all the features. These z-scores thus describe the extremeness of each real SV’s feature relative to their simulations. Subsequently, we applied these normalized features to train an unsupervised model using an active learning approach.

To begin, we developed insertion and deletion models independently. We then randomly withheld SVs in chromosomes 3, 5, 14, 17, 20, and 22 from HGSVC2 during training. We started with all possible flanking, breakpoint, and SV feature annotations to train our model. Subsequently, we reduced the number of features to include only those that displayed statistical significance across any workflow labels based on an ANOVA test. Furthermore, we retained only one of two features if they had correlations exceeding 0.95 to avoid redundancy and to reduce model complexity. We also performed hyperparameter tuning to obtain optimized parameters for our unsupervised clustering algorithm, producing clusters that align but do not strictly require homology thresholds. We chose Hierarchical Density-Based Spatial Clustering of Applications with Noise (HDBSCAN)(59) to maintain the gradient-like relationships observed when comparing HLH, ILH, and NLH, while effectively handling noise and density variations and alleviating the need to select the number of clusters. We optimized various parameters, including the number of dimensions for reduction using PCA (2, 5, 10, 20), the minimum number of points in a cluster (300, 500, 700, 900, 1100, 1300), and the distance metric (Bray-Curtis, Canberra, Chebyshev, Euclidean, Manhattan). We identified “high quality” points assigned to each cluster for every combination of parameters. These points are defined as those with a cumulative confidence higher than 0.5 (i.e., not noise), and those where the difference between the highest and second-highest cluster assignment probability exceeds 0.25 (i.e., SV is confidently assigned to a specific cluster). For each parameter combination we considered, we required that at least 50% of the SVs from each workflow class be classified as “high quality.” In tandem, we assigned clusters a pseudo-label based on the greatest proportion of the workflow label represented (ie. HLH, ILH, NLH, Undefined). Using the workflow labels and their assigned pseudo-labels, we applied Fisher’s Exact tests between the clusters with the greatest proportion of NLH and NLH SVs. Our optimal parameter combination yielded the greatest odds ratio (OR). Together, these requirements ensured large-scale and high-quality clustering while leveraging but not explicitly requiring homology as the sole feature.

Next, we used each clusters’ subset of “high quality” points to train a simple k-nearest neighbor (KNN) model using Euclidean distance. We chose this approach since HGSVC2 SVs are considered the highest quality; therefore, we assumed the patterns underlying these clusters would be the most reliable and discriminative. To find the optimal number of neighbors to consider, we used the following expressions,

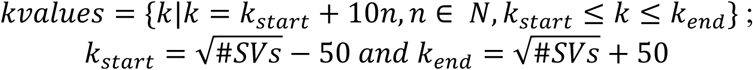

We initialized with 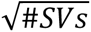 and used a step size of 10, up to 50 (represented by *n*). We conducted 6-fold cross-validation to identify the optimal *kvalue*, using distinct subsets of chromosomes in each fold. Since the mean accuracy remained high across all the folds, we selected the *kvalue* with the least variation in the accuracies. After training an insertion and deletion model, we froze the optimal PCA preprocessing and KNN models, allowing them to be applied to independent datasets without retraining. Following clustering, we considered the z-scores of the alignment lengths. For each cluster, we defined homology-driven (*HD*) and nonhomology-driven (*nHD*) subclusters to discriminate between SVs with z-scores above and below 2, respectively. We then applied these models and subcluster classifications to the held-out HGSVC2 SVs and SVs from 1KG-ONT and 1KG.

### Downstream analyses

Beyond building a homology-based and an unsupervised approach for assigning DSB repair mechanism to label SVs, we performed a series of downstream analyses to characterize the distinct repair classes. For instance, we annotated SV breakpoints using the 18-state H1-hESCs Chrom-HMM(52) annotations to understand epigenetic state differences among various homology-based SV categories. Furthermore, we calculated the standardized residuals (*r*) derived from *χ*_2_ tests to quantify the difference in expected contributions. We also calculated the mean enrichment score (ES) as a complementary approach to quantify enrichment or depletion patterns of SVs overlapping with functional genomic elements, relative to SV simulations. Using the observed and matching simulated SVs, we used bedtools(94) to intersect insertions’ breakpoints or the deleted regions (minimum of 1bp) with several genomic features. As such, SVs were intersected with features including coding regions (CDS), candidate cis-regulatory elements (cCREs) from SCREEN(95) (including proximal and distal enhancer-like signatures), topologically associated domains (TADs)(96), T1 and T2-lamina associated domains (LADs)(97), common and rare fragile sites(98), and regions mapping to >95% Segmental Duplications. We selected this collection of elements to strategically assess the relationships between global negative selection pressures and DSB repair strategies. With these features, we calculated a fraction, then an enrichment score (ES) and empirical p-values;

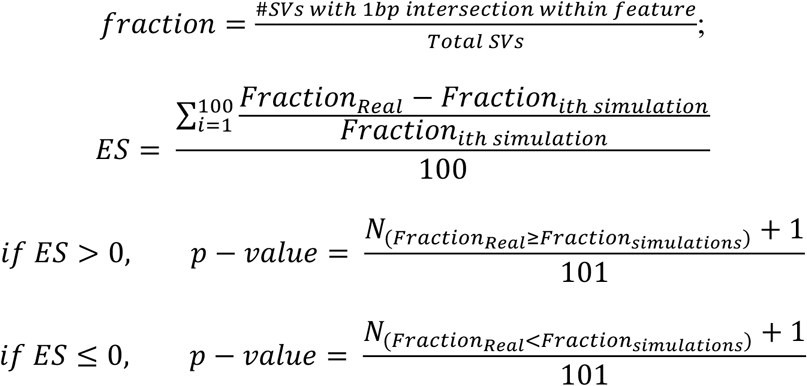

Given that ES is a direct comparison between observed and simulated SVs, an ES>0 represents an enrichment of real SVs within a genomic element. In contrast, an ES<0 represents a depletion pattern. ES was calculated by dataset and label, along with an agglomerated ES (“All together”). We used a similar strategy to assess gene sets, including housekeeping genes and pathways from Reactome(99). In this case, we aimed to probe potential relationships between gene function and DSB repair choice, whereby the selected pathways are alluded to in Ebert et al.(13).

We then used the previously published machine learning model, Orca(61), to investigate the role of repair mechanisms in genomic folding. We filtered for HGSVC2 deletions exceeding 4000bp at a 1Mb resolution, where one pixel represents 4000bp. The results included predicted wildtype (wt) and mutant contact maps, representing pairwise interactions by *log_e_*(*fold over distance based background*). We used the matching wt and mutant 0.5Mb up and downstream contact maps when measuring change. We employed the structural similarity index measure (SSIM) and absolute Spearman’s correlation coefficient as small- and large-scale change-to-folding metrics, respectively(62). SSIM is a measure to describe structural changes as per the perceived similarity of two contact maps. In contrast, Spearman’s correlation can quantify changes in the intensity rank and further be processed to indicate the A/B compartment. We further demonstrated changes in folding by selecting two SVs of similar length and in proximity, chr8-7194444-DEL-8515 and chr8-7283532-DEL-15242. We calculated the difference in 3D interactions by directly comparing their wt and mutant contact maps.

To strategically uncover DSB repair choice inclination in the genome, we investigated SV hotspots by cluster label. We filtered for rare SVs (AF<1%) from the 1KG project to dissociate regional repair bias from common SVs. For each individual, we counted the number of *CI_HD_* and *CII_nHD_* SVs within a 1Mb window across the genome. For each population, we calculated a “hotspot threshold” using the 90^th^ percentile from the SV counts, since baseline variation differs by population. For each population, window, and subcluster class, we then counted the number of individuals who met this SV count threshold, therefore describing individuals with an abundance of SVs in a specific window in a cluster-specific manner. We used the *CI_HD_* and *CII_nHD_* cluster values, to conduct a Fisher’s Exact test, for each population and window. An OR>1 indicates a window enriches *CI_HD_* SVs, while an OR<1 is an enrichment of *CII_nHD_* SVs. Finally, we applied a genome-wide Benjamini/Hochberg p-value adjustment for the tests, at a population level. For visualization, ORs exceeding five were capped to five.

We expect regions with rates of greater homology-driven repair, to display high recombination rates. Therefore, we assigned population-specific recombination rates to the breakpoints of rare SVs from 1KG. For each individual and subcluster class, we calculated the mean recombination rate of their SVs and then determined a cluster and population-specific median recombination rate. To assess population structures, we calculated the absolute difference in median recombination rates for each pair of populations. Upon combining *CI* and *CII* subclasses, when two populations exhibit a large difference in recombination rates for a given class, it indicates greater divergence in the deployment of recombination. For visualization, the populations were organized according to the hierarchical clusters identified by Spence et al(63).

To formally investigate the role of DSB repair choice and pathogenicity, we filtered the 1KG SVs based on those identified in probands. We created three groups of SVs from healthy individuals, which included common (AF > 5%) and rare SVs (AF < 1%) with evidence of inheritance, as well as SVs lacking evidence of inheritance, which are thus considered *de novo* for the proband. Next, we obtained *de novo* SVs from various rare disease cohorts in 100KG using the *de novo* variant dataset, filtering for deletions greater than 50 bp, along with variants that passed the *altreadparent*, *abrasion,* and *patch* filters. This resulted in cohorts of individuals with neurological and neurodevelopmental disorders (NND; n=209), ultra-rare disorders (URD; n=37), cardiovascular disorders (Cardio; n=24), renal and urinary tract disorders (RUT; n=21), endocrine disorders (Endo; n=20), and other disorders (n=85). Following the application of the KNN deletion model and z-score based threshold, we compared the proportion of unique SVs assigned to *CI* and *CII* across these cohorts. Finally, we examined all the z-score based features which underly the cluster labels (ie. *CI* and *CII*) across cohorts. Our goal was to determine whether there was evidence of a feature that could distinguish inherited from *de novo* SVs, as well as SVs from healthy and diseased individuals. We believe this highlighted the importance of studying underlying patterns in repair when considering repair mechanisms, modes of inheritance, and the impact of SVs. To achieve this, we calculated the absolute Cohen’s d effect size across all z-score standardized features, comparing all cohorts and both clustering classes, *CI* and *CII*. From previous work, Cohen’s d values of 0.5 and 0.8 indicate medium and large effect sizes, respectively(58). To evaluate phenotypic differentiation, we focused on participants with URDs, given the notable pattern we observed in local H3K27me3 levels, along with their complex and multisystem phenotypes. We conducted Fisher Exact tests for reported phenotypes by comparing individuals with CI and CII *de novo* SVs, along with the presence or absence (including ‘No’ and ‘Unknown’) of a documented phenotype.

## Results

### Application and validation of local homology-based workflow using several SV catalogs

In our work, we included SVs from two lrWGS (HGSVC(13) and 1KG-ONT(43)) and one srWGS (high-coverage 1KG(44) project) studies to capture germline variant diversity at the population level. Prior research has shown that local homology near breakpoints can influence repair choice(45). Therefore, we extracted genomic regions corresponding to 1975bp upstream or downstream of the SV, along with the first and last 25bp within the SV breakpoint junction. Subsequently, we performed pair-wise sequence alignment between these regions. We only considered alignments with opposing orientations relative to the breakpoints and those that are proximal to the SV breakpoint. Finally, we assessed the alignment length and quality to categorize SVs into distinct homology groups (**Figure 1A**). For example, SVs with high local homology (HLH) required a minimum alignment length of 100 bp and sequence identities above 90%. In contrast, intermediate local homology (ILH) required alignment lengths between 20 and 100bp, with identities greater than 80%. In this context, the ILH class SVs resembled variants induced by SSA and aEJ pathways. While these pathways are functionally distinct, they remain largely understudied and difficult to differentiate using strict homology thresholds(46); therefore, we grouped these SVs together. SVs could also exhibit no local homology (NLH) or be categorized as “undefined” if their length or alignment quality did not meet these thresholds. Our homology length and quality thresholds reflect experimentally derived values from proteins involved in HR, SSA, aEJ, or NHEJ(46–49). Upon applying this workflow, we found that NLH represents the largest proportion of SVs, regardless of the SV dataset (**Figure 1B; Figure S1**). For instance, we observed that NLH, ILH, HLH, and undefined classifications accounted for 52.6%, 26.0%, 7.6%, and 13.8% of unique insertions from HGSVC2, respectively. We found similar contribution patterns for 1KG-ONT insertions, classified as NLH (47.3%), ILH (27.1%), and HLH (5.3%) by our homology-based workflow. These proportions remained consistent among deletions. We found that the distribution of SV lengths among NLH, ILH, and HLH SVs was similar; however, Undefined SVs skewed toward longer length values **(Figure S2)**.

**Figure 1.**
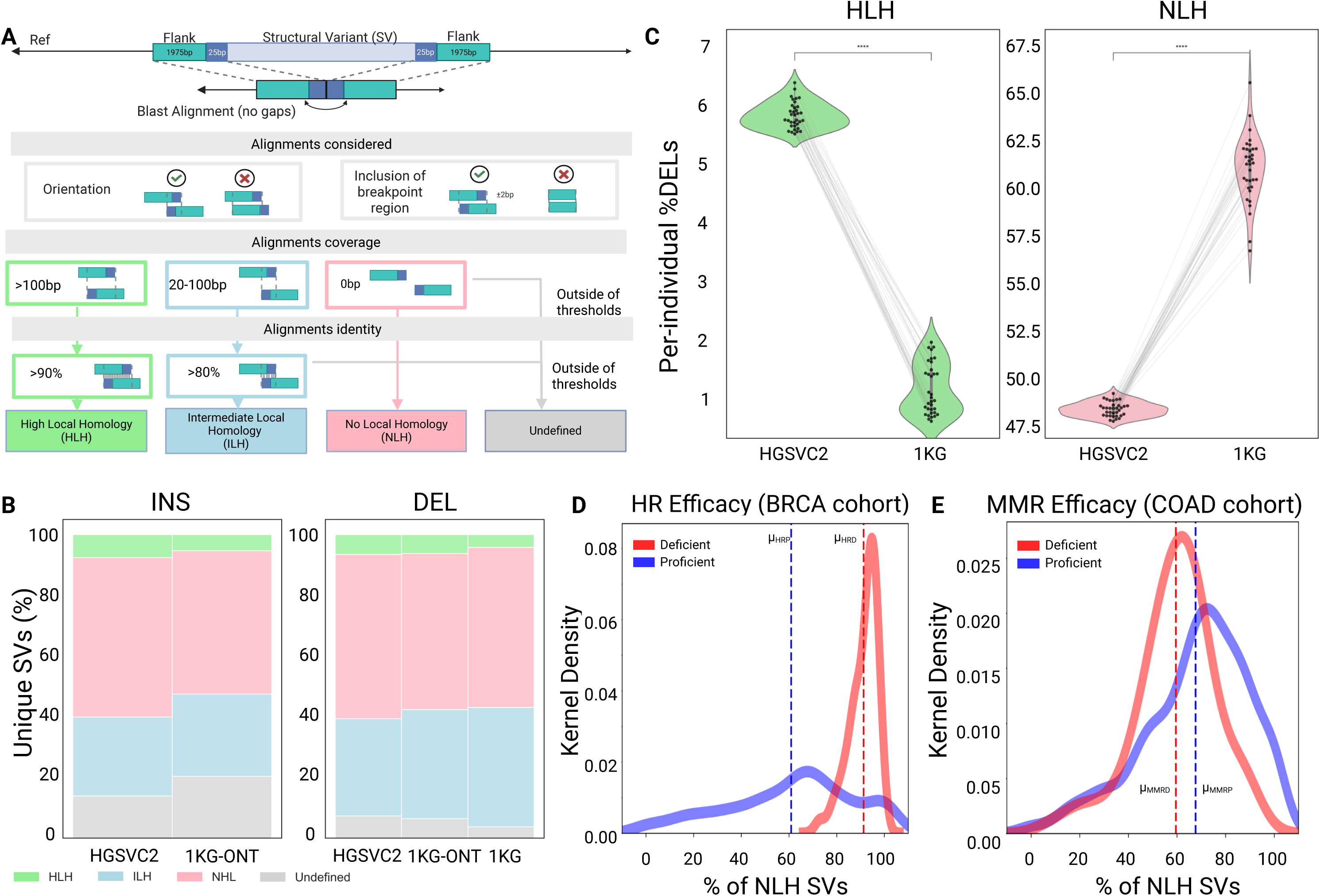
The homology-based framework to describe SV introduction as per DSB repair mechanisms. **(A).** Schematic of the homology-based label workflow. The 1975 bp upstream and downstream sequences local to deletions (depicted) and insertions are aligned with 25bp of the SV without gaps using BLAST. Alignment orientation and the inclusion of the breakpoint are assessed. Quality and length determine the SV class. **(B).** Proportion of workflow labels applied to unique SVs in lrWGS (HGSVC2 & 1KG-ONT) and srWGS (high coverage 1KG phase 3) datasets. **(C).** Per-individual proportion of deletions labeled with the workflow when applied to individuals in both HGSVC2 and 1KG (n=34). The proportions of individuals are connected with grey lines in the violin plots. (Paired t-tests). **(D-E).** The distribution of resolved somatic deletions (50 bp+) labeled as HLH among the 100KG project cohorts, visualized with kernel density estimate (KDE) plots. The selected cohorts demonstrate distinct patterns of repair to validate the workflow. The breast cancer (BRCA) group served as the positive control, to monitor HR efficacy **(D),** while the colorectal cancer (COAD) cohort served as the negative control, to observe MMR **(E).** Repair-deficient cohorts are shown in red, while repair-proficient cohorts are in blue. Dotted lines depict a cohort’s mean %HLH.

We subsequently utilized the SVs from the HGSVC2 project to compare our results to another sequence-based SV mechanism tool, Breakseq2(25). Although Breakseq employs distinct homology metrics and incorporates additional repeat annotations, we observed a general alignment between the two methods. We found that 89.9% of HLH-labeled insertions were classified as NAHR, while 7.6% were classified as non-homologous recombination (NHR; **Figure S3A**). This pattern also applied to deletions, with 74.3% and 2.2% of HLH SVs categorized as NAHR and NHR, respectively (**Figure S3B**).

We found minor differences in the label assignment proportions when analyzing the lrWGS dataset (HGSVC2) and the srWGS dataset (1KG project). Therefore, we compared these datasets using SVs at an individual level, using individuals common to both studies (n=34). We assumed HGSVC2 SVs are of the highest quality, as they were identified from haplotype-specific assemblies derived from lrWGS (PacBio) sequencing data. As expected, we found that among deletions, the per-individual proportion of HLH SVs in HGSVC2 (*μ*_*HGSVC2*_ = 5.85%) was significantly higher than 1KG (*μ_1KG_* = 1.11%; p-value= 1.59e-36; **Figure 1C**). In contrast, the NLH proportions per individual were significantly lower for HGSVC2 (*μ_HGSVC3_* = 48.5%) compared to the 1KG dataset (*μ_1KG_* = 60.87%; p-value= 1.58e-29).

To validate our overtly simple homology-based workflow, we utilized SVs from samples with known extreme repair patterns. Therefore, we constructed sample cohorts of short-read somatic SVs from the 100,000 Genomes (100KG)(50) project to demonstrate changes or the absence of change in the proportion of NLH SVs. We used breast and colorectal cancer samples (BRCA; COAD) to leverage HR and mismatch repair (MMR) proficiency, respectively(51). We found that HR-deficient samples tend to have more NLH deletions (91.2%) than HR-proficient samples (60.8%; p-value = 2.3e-29; **Figure 1D**). In contrast, although still statistically different (p-value = 1.2e-08), the MMR-deficient (59.5%) and proficient samples (67.5%) exhibited a much smaller difference in NLH assignment (**Figure 1E**). These findings illustrate how the workflow uniquely captures SVs introduced by homology-based repair mechanisms.

### Genomic elements are uniquely enriched among homology-based workflow labels

Our homology-based workflow connects DSB repair mechanism patterns with SVs. However, our simple workflow does not consider additional sequence and epigenetic features that likely influence DSB repair choice. These patterns are particularly interesting since previous studies exploring epigenetic differences among DSB repair strategies have revealed inconsistencies regarding whether certain DSBs in heterochromatin regions tend to be repaired via HR, aEJ, or NHEJ. Variations in experimental models and their inherent biases may explain these discrepancies(34). Therefore, we utilized our extensive catalog of genome-wide SVs to better understand global genomic and epigenomic patterns in repair. To accomplish this goal, we applied the 18-state ChromHMM(52) annotations from H1-hESC (H1) cell lines as a proxy for the germline(53). We found that upstream breakpoints of HLH deletions favor heterochromatin regions (standardized residual (*r*) *r_HLH_* = 6.5; **Supplement Table 1A**, **Figure 2A**), while ILH breakpoints overlap with weak (*r_ILH_* = 11.7) and strong transcription sites (*r_ILH_* = 7.4). Furthermore, NLH breakpoints predominantly occupy epigenetically quiescent regions (*r_NLH_* = 10.6) within the genome. We observed a similar pattern of association between chromatin states and homology profiles for flanking regions downstream to breakpoints in deletions and insertions **(Supplement Table 1B; Figure S4)**.

**Figure 2.**
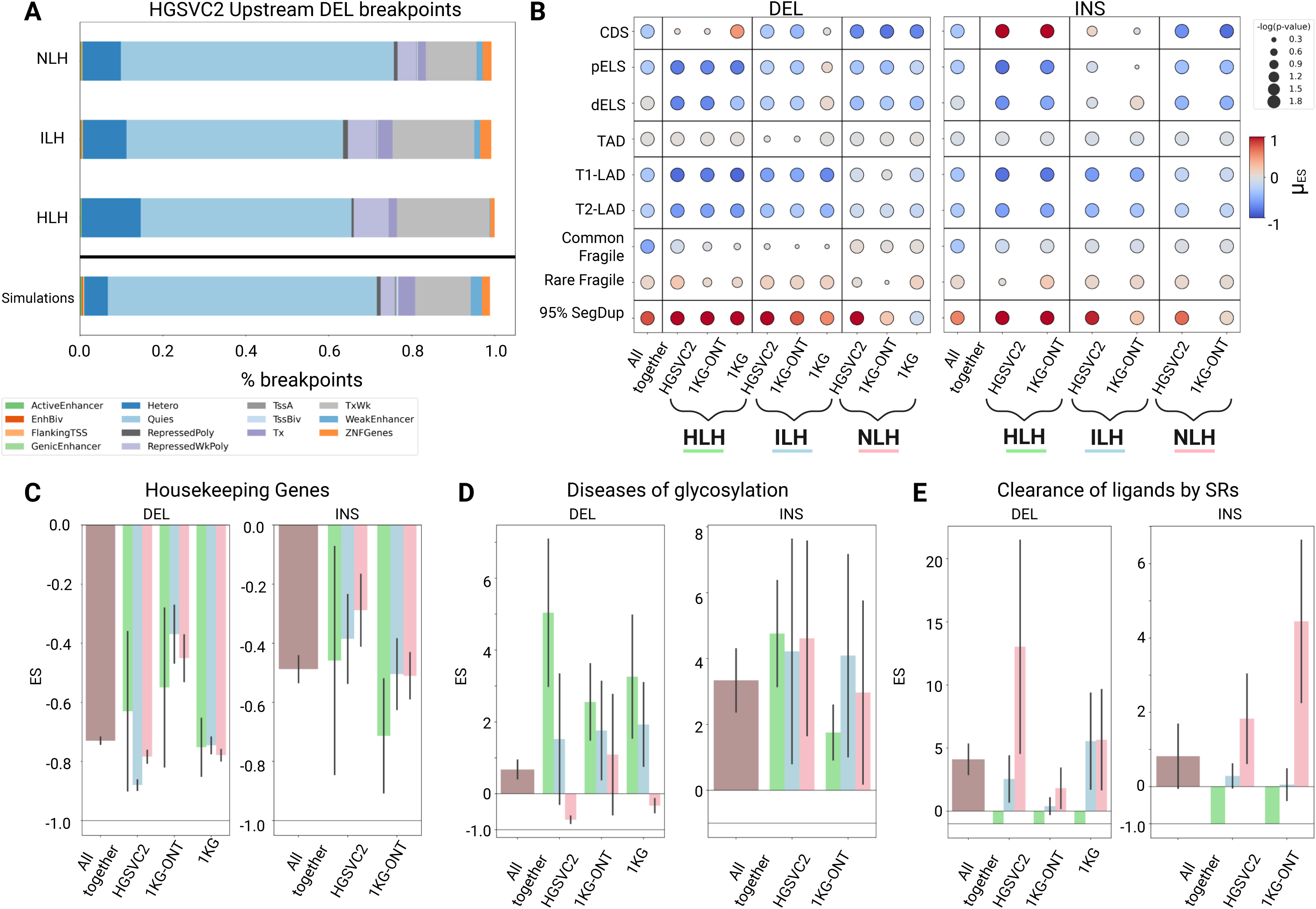
Enrichment of genomic elements, highlighted as per the homology-based workflow. **(A).** Stacked bar plots illustrate the proportion of upstream deletion breakpoints from HGSVC2 that fall within ChromHMM states. The bottommost bar represents these proportions across all simulations, while the bars above show the breakdown by label (HLH, ILH, NLH). **(B).** Enrichment and depletion patterns of SVs across genomic elements are depicted. Each marker indicates the magnitude, direction, and statistical significance of the mean enrichment score (ES). Magnitude and direction are differentiated by color, whereas statistical significance (represented as -log(empirical p-value)) is indicated by the marker size. The leftmost column demonstrates the aggregated ES among deletions and insertions, while the other columns are specific to workflow labels and datasets. The elements considered include coding regions (CDS), proximal and distal enhancer-like elements (pELS, dELS), topologically associated domains (TADs), T1 and T2 lamina-associated domains (LADs), common and rare fragile sites, and segmental duplications above 95% sequence identity. For visualization, the ES color intensity was capped between -1 and 1. **(C-E).** Bar plots illustrate the ES for deletions and insertions subject to CDS perturbation in gene sets. Mean ES (bar height and direction) and standard deviations (error bars) reveal the aggregated enrichment patterns, along with label and dataset specifics. Gene sets include housekeeping genes (HRT Atlas v1.0; **C**), Diseases of glycosylation (R-HSA-3781865; **D**), and binding and uptake of ligands by scavenger receptors (SRs; Reactome; R-HSA-2173782; **E**).

Beyond chromatin state differences, we examined the variability in selection pressures observed among distinct categories of SVs (based on the simple workflow classes) affecting various genomic elements. We generated 100 simulated SVs for each observed SV using StructURal Variant Majority Vote (SURVIVOR)(54) tool to quantify these selection pressures. Briefly, SURVIVOR utilizes a specified length, class, and reference genome to generate simulated SVs that match in length, SV type, and chromosome. We then calculated a mean enrichment score (ES) using the expected and observed SV profiles. Overall, we observed significant depletions of SVs overlapping with various functional elements in the genome (**Figure 2B**), regardless of the homology-based label, aligning with prior studies(55). However, we found variability in the magnitude and direction of these selection pressures among distinct SV categories. For example, among candidate cis-regulatory elements (cCREs) such as proximal enhancer-like signatures (pELS), the depletion strength for deletions in HGSVC2 is much stronger among HLH SVs (ES_HLH_=-0.779, p-value=0.009) compared to ILH (ES_ILH_=-0.246, p-value=0.009) and NLH SVs (ES_NLH_=-0.345, p-value=0.009). This pattern was generally consistent across other genomic elements, including distal enhancer-like signatures (dELS), lamina-associated domains (LADs), and rare fragile sites. Interestingly, HLH insertions are enriched in segmental duplications (SD; ES_HLH_=1.89, p-value=0.009). However, for deletions, there was no significant enrichment/depletion pattern in HLH SVs at CDSs (ES_HLH_=0.08, p-value=0.326). Therefore, we found that the workflow-based labels align with paradoxes in the literature, where homology-based repair appears functional in both heterochromatin and euchromatin regions.

Considering the surprising enrichment pattern of HLH insertions overlapping with CDS elements, we investigated whether a gene’s function could be driving this observation. Therefore, we quantified the enrichment score of SVs overlapping with specific gene sets, selected based on the original HGSVC2 study(13). First, we quantified enrichment pattern of SVs affecting housekeeping genes and expectedly found negative selection pressure, irrespective of the SV class (HGSVC deletions: ES_HLH_= -0.629; ES_ILH_= -0.879; ES_NLH_= -0.783; **Figure 2C**). In contrast, we observed that HLH and ILH deletions are especially enriched in the diseases of glycosylation gene set (HGSVC deletions: ES_HLH_= 5.03; ES_ILH_= 1.52; ES_NLH_= -0.721; **Figure 2D**), while ILH and NLH SVs are more likely to impact genes in the scavenger receptor binding and uptake pathway (HGSVC deletions: ES_HLH_= -1.00; ES_ILH_= 2.55; ES_NLH_= 13.03; **Figure 2E**). However, when considering insertions at these receptor binding genes, the strength of this enrichment was weaker and less divergent for NLH SVs (HGSVC insertions: ES_HLH_= -1.00; ES_ILH_= 0.291; ES_NLH_= 1.83). While these results seek to connect repair mechanisms to gene function, the presence or absence of local homology may be an inherent characteristic of the surveyed genes. For instance, mucins in the glycosylation gene set are filled with variable number tandem repeats(56), while the variation in immune-related genes can be introduced via NHEJ(57). Therefore, for these gene sets, there may be a built-in increase in the enrichment of HLH and NLH SVs, respectively.

### DNA shape and epigenomics-associated features can discriminate between workflow labels

Our observations of differences in chromatin states and selection patterns for distinct homology-based SV classes indicate the role of additional genetic and epigenetic features that may facilitate DNA repair. Thus, we comprehensively characterized the features that underlie SVs, including sequence attributes, DNA shape, repetitive motifs, epigenetics (histone profile and DNA accessibility), and replication timing. We calculated the Z-score of these features for upstream and downstream flanks, breakpoints, and within SV regions using the observed and simulated datasets (**Supplement Table 2**, **Figure 3A**). We obtained 135 and 109 features for deletions and insertions, respectively. Using the HGSVC2 dataset and workflow-based labels, we employed Cohen’s d effect sizes for pairwise comparisons (HLH vs. ILH, HLH vs. NLH, and ILH vs. NLH) to identify features contributing to their discrimination. Cohen’s d measures the standardized difference between the means of two groups, where effect sizes of 0.5 and 0.8 are considered medium and large effects, respectively(58). We observed small effect sizes for most features when comparing HLH to ILH (Figure 3B & S5). However, there was greater separation between HLH and NLH SVs for features related to sequence, with a mean Cohen’s d of 0.623. Closer examination revealed that the top 15 most discriminatory features between HLH and NLH deletions included %GC content, DNA methylation, local electrostatic potential (EP), and minor groove width (MGW; Figure 3C). We visualized the average methylation of upstream flanking sequences and local MGW variation in deletions using homology-based labels (Figure 3D-E). Notably, HLH SVs showed high normalized mean methylation and low normalized MGW variation. Additionally, the z-scores for NLH SVs were near 0, indicating similarity between observed and simulated SVs, suggesting a background repair mechanism. Furthermore, for most features, including methylation and MGW, undefined SVs had z-scores similar to NLH SVs. These findings were generally consistent across insertions (**Figures S6-7**). Altogether, our analyses suggest that a broader inventory of features may be utilized to differentiate between homology-based labels, thereby providing evidence that these features may support DSB repair.

**Figure 3.**
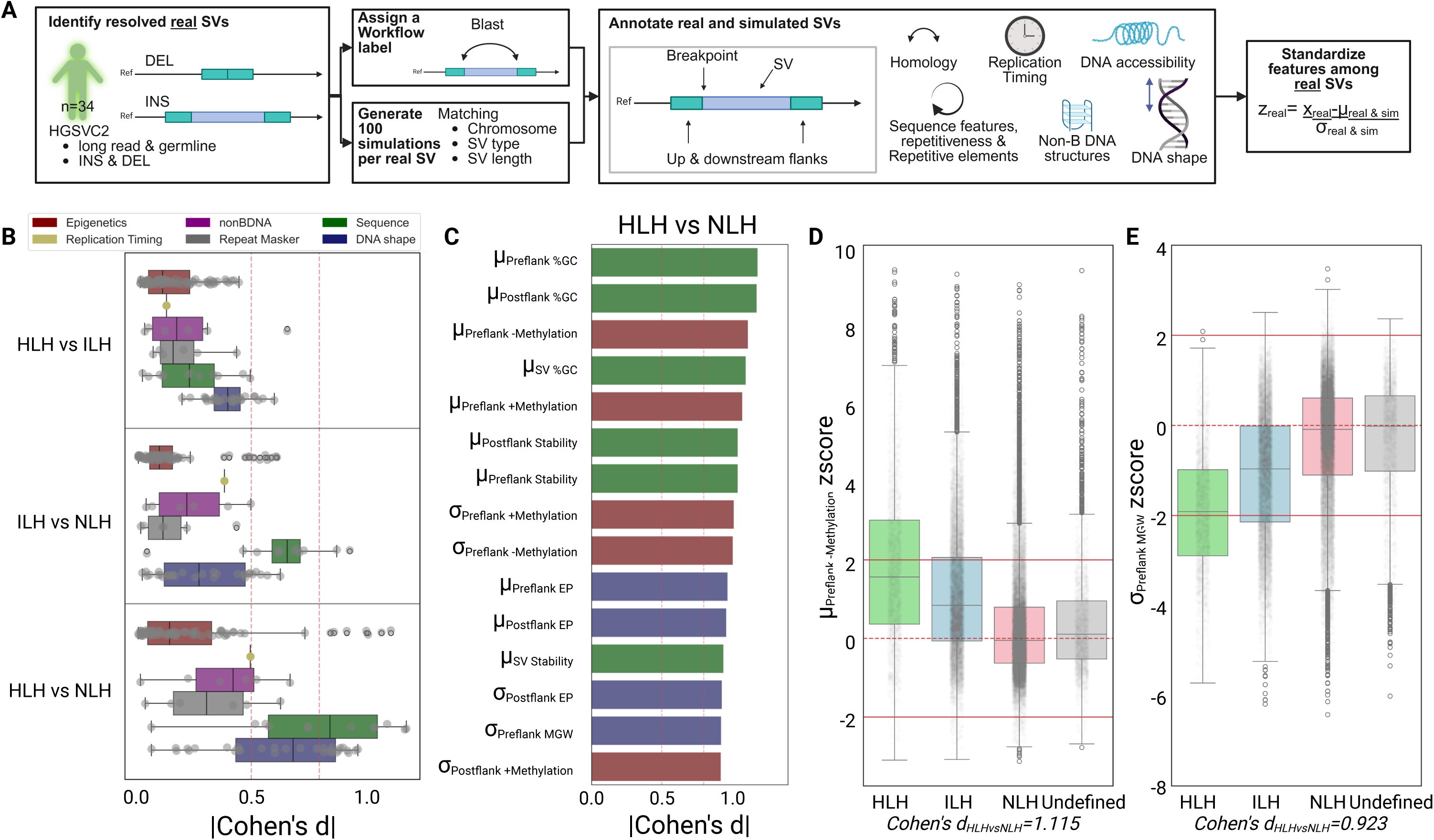
Probe into underlying features which discriminate homology-based labels. **(A).** A schematic depicting the strategy to standardize SV, with 2000bp flanks and breakpoint features. HGSVC2 SVs are labeled with homology-based annotations. In tandem, 100 simulated SVs are generated using SURVIVOR for each SV. These simulated SVs match the observed SV’s chromosome, SV type, and SV length. Each observed and simulated SV is then annotated with features describing their local homology, sequence (including repetitive elements), replication timing, presence of non-B DNA motifs, epigenomics (derived from ChIP-Seq, DNA methylation, and DNAase-seq), and DNA shape. Finally, a standardized z-score is calculated for each feature. (B-E). Underlying patterns supporting HGSVC2 deletions. **(B).** Boxplots depicting the absolute effect sizes of deletions. Each boxplot illustrates a pairwise comparison between two workflow labels for a feature type, with effect sizes measured using Cohen’s d. Medium and large effect sizes are indicated with red dotted lines at 0.5 and 0.8, respectively. **(C).** A barplot highlighting the top 15 most discriminatory features based on effect sizes between HLH and NLH. The color of the bar corresponds to the feature type. **(D-E).** Boxplots illustrating z-scores by workflow label according to the mean level of upstream DNA methylation **(D)** and the variation in upstream minor groove width (MGW; **E).** For SVs with z-scores close to 0, the distributions of the observed and simulated SVs for the given feature are similar, as depicted by a red dotted line. Extreme z-scores, those above 2 or below -2, are shown with solid red lines.

### Harnessing discriminatory features beyond homology to infer the mechanistic origin of SVs

Given that additional features beyond local sequence homology may describe patterns in repair choice, we trained an active-learning-based unsupervised model to classify SVs (**Figure 4A**). We reasoned that an unsupervised approach was most appropriate, as strict homology thresholds might be prohibitive for capturing the complete landscape of features that could contribute to an SV’s introduction. Nonetheless, strict homology-based high-confidence data points can serve as pseudo-labels to guide the unsupervised model toward our biological expectations, thereby emphasizing the importance of local homology. In this approach, we first used hierarchical density-based spatial clustering of applications with noise (HDBSCAN)(59) to perform high-quality SV cluster assignments. Subsequently, we quantified cluster enrichment in NLH and HLH SVs using the odds ratio (OR) and required that a substantial proportion of SVs be represented in the resulting clusters. Several models were evaluated by altering parameter combinations for feature dimension reduction, cluster size, and distance metrics. We then selected the model that reflected the highest OR, thereby identifying clusters with the most significant enrichment of NLH and HLH. Next, the high-quality SVs and their pseudo-label assignments were passed on to k-nearest neighbors (KNN) models. We trained separate models for insertions and deletions and utilized randomly held-out chromosomes (chr 3, 5, 14, 17, 20, 22) for model evaluation. Thus, our method employs additional underlying features while leveraging the original workflow-based (ie. HLH, ILH, NLH) labels without imposing thresholds.

**Figure 4.**
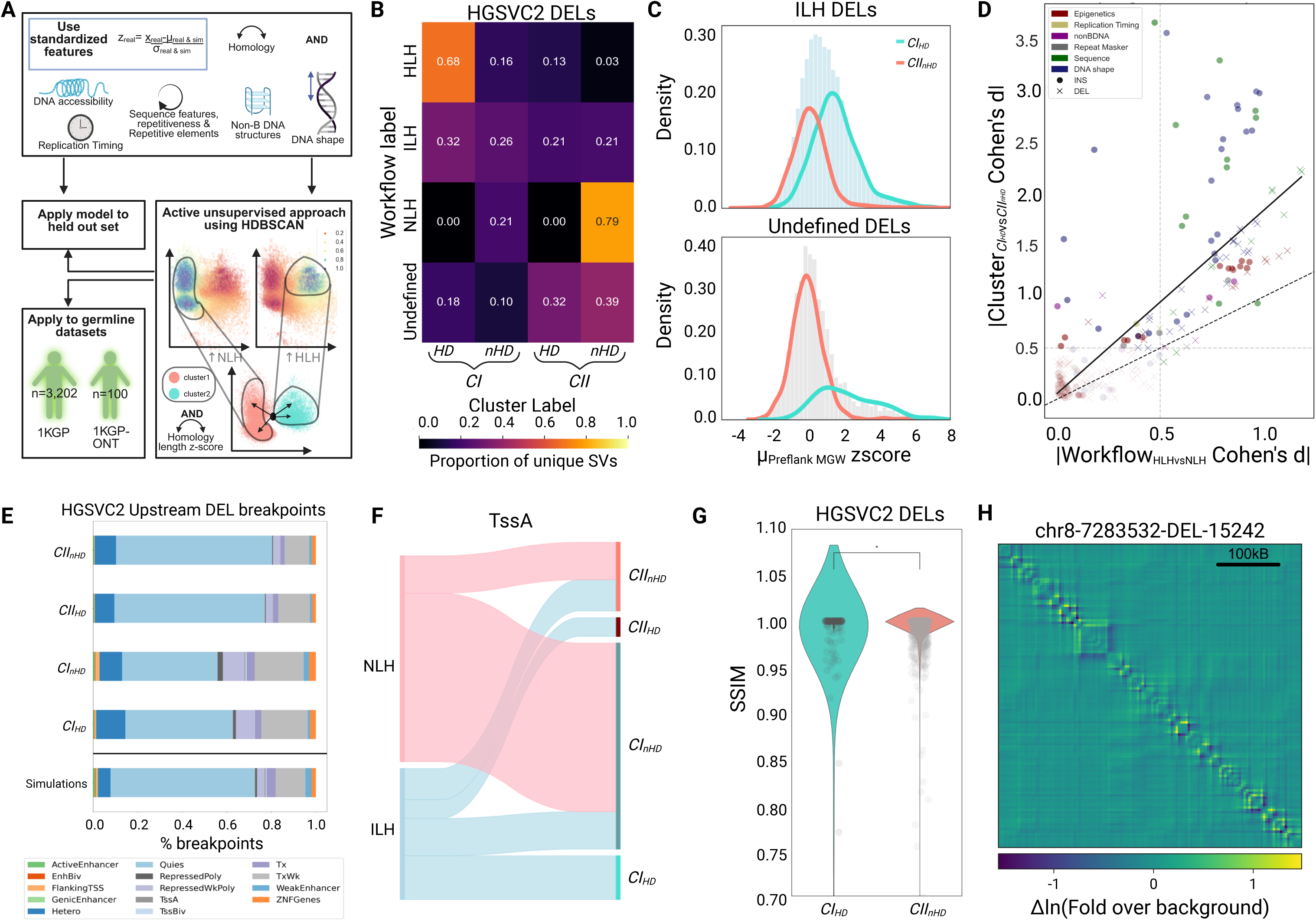
Active unsupervised clustering approach which exploits but does not strictly require thresholds for local homology. **(A).** Graphic of the active unsupervised clustering approach used to exploit features underlying SVs. SVs from HGSVC2 in chromosomes 3, 5, 14, 17, 20, 22 are held out. An insertion and deletion model were independently developed. The z-score standardized features are filtered. Upon optimizing preprocessing and parameters for Hierarchical Density-Based Spatial Clustering of Applications with Noise (HDBSCAN), clusters representing an enrichment in HLH and NLH SVs are identified. SVs with the highest quality cluster assignment were used to train a k-nearest neighbour (KNN) model. The models were frozen and applied to the held out HGSVC2 SVs and SVs in 1KG and 1KG-ONT. Subsequently, SVs with z-scores above 2 in alignment length were labeled as homology driven (HD), while others were labelled nonhomology driven (nHD). **(B).** Heatmap visualizing the concordance between the homology-based and subclustering-based labels among the HGSVC2 deletions. Proportions are row-wise. **(C).** Histograms and KDE plots depicting the distribution of z-scores from the upstream mean minor groove width (MGW). The top plot depicts HGSVC2 deletions labelled as ILH, while the bottom plot contains Undefined deletions. On both plots, and for both workflow-based labels, KDE plots show the distribution of z-scores among the subcluster-based labels, *CI_HD_* and *CII_nHD_*. **(D).** scatterplot comparing the effect sizes based on the workflow-based label (HLH versus NLH) and the cluster-based labels (*CI_HD_* versus *CII_nHD_*). A linear regression was fitted after excluding the homology-based features and is shown with a solid black line. The dotted y=x line serves as a comparison, demonstrating identical performance. Features with consistently low effect sizes, falling below 0.5 in both labeling strategies, are shown in grey. The remaining points are colored by feature type, and the shape of the markers corresponds to SV type. **(E)** Stacked bar plot depicting the breakpoints of upstream breakpoints from deletions falling within ChromHMM states according to the cluster-based labels. **(F)** Sankey plot illustrating insertions at ChromHMM’s active transcription sites (TssA). The left side displays the homology-based label, while the right side corresponds to the cluster-based label. **(G)** Boxplots comparing the local changes in genomic folding based on the structural similarity index measure (SSIM) between *CI_HD_* and *CII_nHD_* SVs (Mann-Whitney). **(H)** The change in local DNA interactions with the *CI_HD_* SV, chr8-7283532-DEL-15242. The 500 Mb wildtype and mutant windows downstream of the SV were directly compared.

Optimal clustering for deletions involved reducing feature dimensions to five components, using Bray-Curtis distance, and a minimum of 700 SVs per cluster (OR_HLH vs NLH_ = 125.59; **Supplement Table 3A)**. This approach utilized 121 features (**Supplement Table 4A)** to generate two clusters. For insertions, we reduced the components to two, used the Canberra distance, and required 1,300 SVs per cluster (OR_HLH vs NLH_ = 54.41; **Supplement Table 3B),** resulting in four clusters based on 95 features (**Supplement Table 4B)**. These clusters were then assigned pseudo-identities to train and validate a KNN model. The optimized KNN models for insertions and deletions used 118 and 96 neighbors, respectively. Next, we sub-categorized SVs within each cluster into homology-driven (HD) and nonhomology-driven (nHD) based on high and low normalized homology-based alignment length scores, respectively. Overall, we generated four and eight sub-clusters for deletions and insertions, respectively.

The trained KNN models and normalized alignment length thresholding were subsequently applied to the held-out SVs from HGSVC2 (Figure S8), 1KG-ONT (Figure S9), and 1KG (Figure S10) datasets. We calculated the concordance between the homology-based workflow and clustering labels using HGSVC2 SVs (Figure 4B & S11). Among the deletions, we identified two clusters (CI, CII), and for insertions, we identified four clusters (CI-IV). In order, CI, CII, CIII, and CIV had the highest proportions of HLH, NLH, ILH, and Undefined SVs, respectively. Furthermore, we subdivided these four clusters into HD and nHD subcategories/sub-clusters based on the homology criterion. This method allowed us to maintain rules regarding the expectation of homology in repair (Figure S12) while capturing underlying patterns. For example, the HLH and CI groups both show high standardized mean upstream flanking %GC, whereas the NLH and CII groups show the opposite (Figure S13). Overall, we observed strong agreement between the workflow and clustering-based methods. For instance, our approach labeled 83.9% of HLH deletions and 78.9% of NLH deletions as CI and CII, respectively. Interestingly, most undefined deletions were assigned to (*CII_HD_* at 32.4%, & *CII_nHD_* at 39.3%).

Next, we examined specific cases where clustering improved the characterization of SVs compared to homology-based labeling. For example, the ILH deletions, chr1-248746661-DEL-59 and chr1-246387882-DEL-312, are in proximity to each other, with equivalent alignment length and quality (**Figure S14AB)**. However, chr1-248746661-DEL-59 was assigned to *CI*, while chr1-246387882-DEL-312 was assigned to *CII.* A close inspection of their local contexts suggested that chr1-248746661-DEL-59 is nested within a direct repeat (DR) with high %GC and consistent MGWs. In contrast, only the downstream flank of chr1-246387882-DEL-312 falls within a DR in a low %GC region with fluctuating MGWs (**Figure S14CD**). This indicates that although these SVs share significant homology at their breakpoint junctions through clustering, their local contexts may also influence the differentiation of underlying repair mechanisms. Similarly, to explore other ambiguous cases, we analyzed the clustering of ILH and Undefined deletions. First, we examined the distribution of features such as the mean MGW of the upstream flank. Previous studies indicate that MRX/N, the complex that initiates DSB processing, can bind at the minor groove(60), and even favors DNA sequences with greater local MGW(40). We observed that, expectedly, the local mean of MGW z-scores from HLH (*μ*_JBK_ = 1.94), ILH (*μ*_JBK_ = 1.03), and NLH (*μ*_JBK_ = 0.75) deletions decreases. We observed that following sub-clustering, ILH (*CI_HD_* = 1.52, *CII_nHD_* = 0.08), and Undefined (*CI_HD_* = 2.14, *CII_nHD_* = −0.11) deletions could be further partitioned using mean MGW z-scores (**Figure 4C).** This was particularly evident, as the distribution of Undefined SVs MGW z-scores has a long right tail, which is subsequently labelled to the *CI* cluster. To confirm the generalizability of these improvements, we directly compared the effect sizes between HLH and NLH to those between the *CI_HD_* and *CII_nHD_* subclusters (**Figure 4D**). Although some features were maintained or even decreased in their ability to differentiate the classes, excluding homology-based features resulted in a significant performance improvement (β = 1.787, p-value = 1.878e-49, R^²^ = 0.601). We found that the clustering-based strategy enhanced the effect sizes of several features, including sequence, DNA shape, replication timing, and epigenetics, specifically for DNA methylation, chromatin openness (as measured by DNase-seq), and H3K27me3 features.

Subsequently, we recharacterized ChromHMM states using our clustering labels (**Figure 4E; Figure S15; Supplement Table 3AB**). We found that the upstream breakpoints of *CI_HD_* deletions are enriched in weak transcription regions 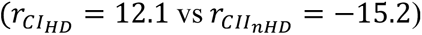 and weak repressed polycomb sites 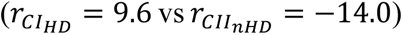. In contrast, *CII_nHD_* SVs displayed very strong enrichment in quiescent regions 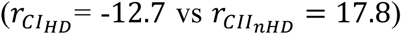. Most interestingly, we found that bivalent transcription start sites 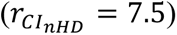 were uniquely enriched among *CI_nHD_* SVs. We then examined the ChromHMM state of SVs, demonstrating inconsistency between the clustering and workflow methods. For instance, we found that NLH SVs in active transcription start sites (**Figure 4F**) and those in repressed polycomb complex regions (**Figure S16**) were primarily assigned to *CI*. These findings indicate that the clustering-based strategy learns features beyond local homology when assigning cluster identities.

Given the striking enrichment of *CII* SVs in quiescent regions, we hypothesized that the breakpoints of *CII* SVs in these areas would cause minimal changes to 3D genomic folding. We employed a deep learning method, Orca(61), which predicts 3D genome contact maps based on genomic sequences. We generated mutant and wild-type contact maps using HGSVC2 deletions of length 4 Kbp or greater. Subsequently, we calculated the absolute correlation and the structural similarity index measure (SSIM) to quantify global and local folding perturbations, respectively(62). We found no difference in global folding changes when comparing the effect of *CI_HD_* and *CII_nHD_* SVs on the 3D-genome organization (**Figure S17A**). However, we observed significant local changes (p-value = 0.0279; **Figure 4G; Figure S17B**). We visualized the interaction differences between the mutant and wild-type maps to demonstrate these changes. We used a 0.5 Mbp downstream window from an *CI_HD_* (chr8-7283532-DEL-15242; **Figure 4H**) and *CII_nHD_* (chr8-7194444-DEL-8515; **Figure S17C-D**) SV. These SVs were selected due to their similar lengths and proximity. Although we found no global rewiring in the 3D structure due to the SV, the *CI* SV introduces small-scale changes across the entire window. Together, this suggests a potential relationship between repair choices and alterations to 3D-genome organization.

### Applications to population structure & *de novo* SVs from rare disease cohorts

We applied our unsupervised method to characterize and compare SVs with distinct underlying repair patterns across diverse populations. Given the vast number of ancestries included, we utilized deletions from the 1KG study for this analysis. We hypothesized that SV (*CI_HD_* and *CII_nHD_*) hotspots, shared across ancestries, may indicate regional biases for a particular repair mechanism, thereby explaining patterns of variation in this region. Since we did not expect significant deviations among different ancestry groups for common SVs, we only included rare SVs (Allele Frequency (AF) < 1%) in this analysis. To identify these hotspots, we divided each chromosome into non-overlapping 1 Mb windows for each individual and calculated the total number of deletions annotated as *CI_HD_* or *CII_nHD_* SVs. We then computed the 90^th^ percentile for these counts within each ancestry group, using this as a threshold to determine how many individuals exceed this SV count for a given window. We quantified a window’s association with the *CI_HD_* or *CII_nHD_* clusters by applying Fisher’s Exact test. We found very few consistent *CII_nHD_* windows across all ancestries (**Figure 5A, Figure S18-19**). However, we observed many consistent *CI_HD_* hotspots, typically located in sub-telomeric regions, gene-dense areas, and within segmental duplications, largely coinciding with regions of strong transcription.

**Figure 5.**
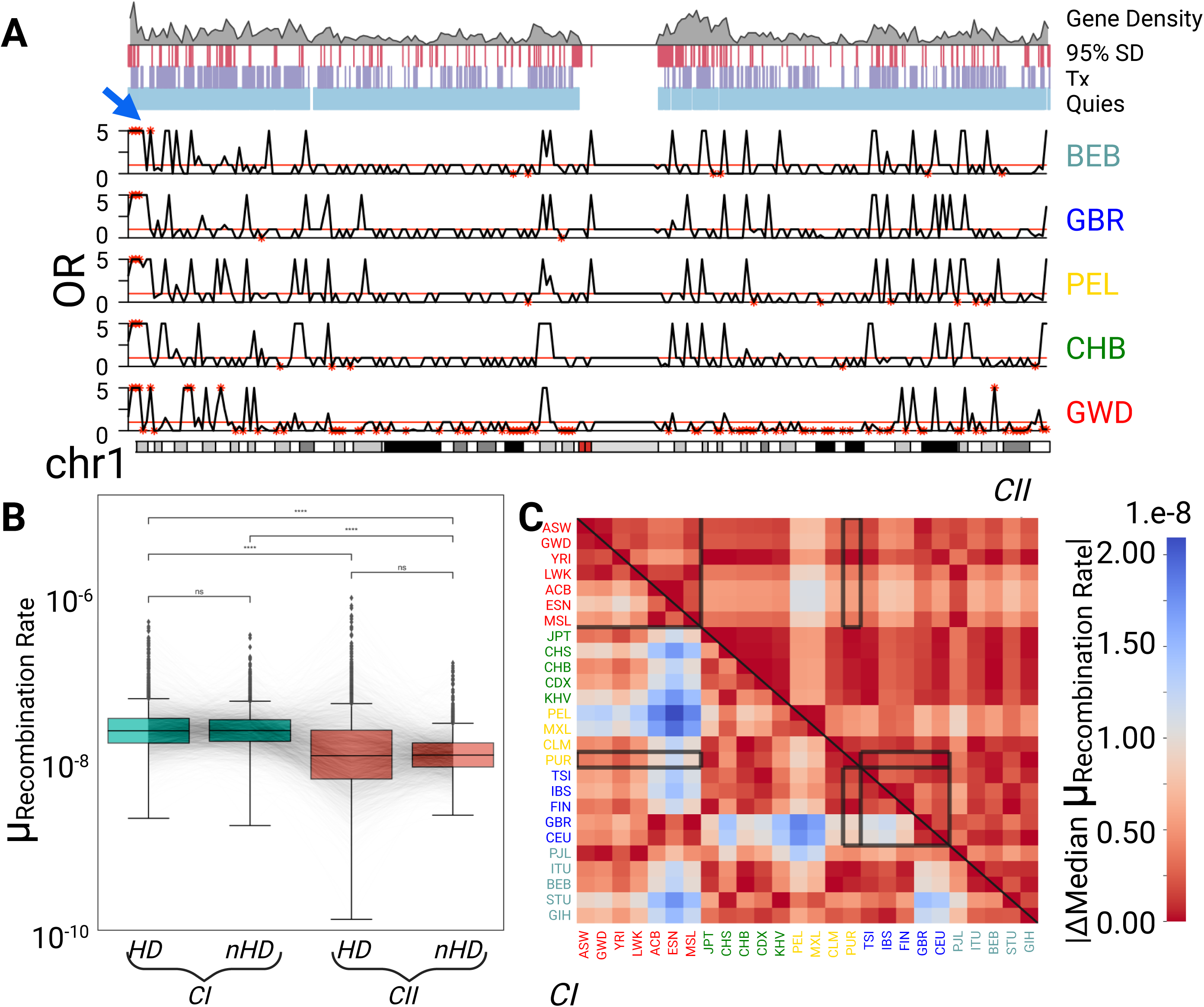
Application of clustering-based strategy to explore ancestry. **(A).** Karyoplot of SV hotspots across 1 Mb windows among rare deletions from 1KG on chromosome 1. The chromosome’s ideogram is displayed at the bottom. Above this, using one population from each superpopulation, the odds ratio illustrating the enrichment of *CI_HD_* SVs is indicated with black lines. A thin red line marks the level at 1, showing no enrichment in *CI_HD_* or *CII_nHD_* SVs in the window. Furthermore, windows with statistical significance, after p-value adjustment, are represented with red stars. A blue arrow points to windows showing consistent and significant hotspots across ancestries. Above these plots are genomic elements aligned to the chromosome. These elements include gene density, 95% segmental duplications (SD), as well as strong transcription (Tx) and quiescent (Quies) sites from ChromHMM. The populations included Bengali in Bangladesh (BEB) from South Asia, British in England and Scotland (GBR) from Europe, Peruvians from Lima (PEL) classified as admixed American, Han Chinese in Beijing (CHB) from East Asia, and the Gambians in Western Division-Mandinka (GWD) from Africa. Text colors correspond to superpopulation. **(B).** Box plot comparing an individual’s mean population-specific recombination rate among the four subclusters. Individuals’ mean rates are connected by grey lines in the boxplots. (Paired t-tests). **(C).** Heatmap illustrating the magnitude of the pairwise differences in recombination rate. For each population, the median of the individual-level mean recombination rates was calculated for *CI* and *CII* SVs. For each population pair, the difference is visualized. The upper and lower triangles display the *CII* and *CI* differences, respectively. Black lines in both the upper and lower triangles highlight the east-west African clines, Puerto Rican admixture, and the north-south European cline.

Since it is well established that sub-telomeric regions are hotspots for recombination(63, 64), and we did not use recombination rate as a feature during clustering, we confirmed significantly higher recombination rates amongst *CI_HD_* (*μ* = 3.65*e* − 0.08) compared to *CII_nHD_* SVs (*μ* = 1.68*e* − 0.08; p-value = 1.06e-147 **Figure 5B**). Furthermore, we investigated recent population-specific dynamics in variation, such as geographical clines and migratory patterns, by utilizing the 1KG population information(63). We found no significant difference in recombination rates within subclusters **(Figure 5B)**; therefore, we combined the *HD* and *nHD* subclusters to investigate population-specific differences among *CI* and *CII* SVs. We found pairwise differences in recombination rates by cluster after calculating the median of a population’s mean recombination rate. We visualized these differences using a previously determined population structure(63) and observed that population dynamics were pronounced only in *CI* (**Figure 5C**). Most notably, the recombination rates of genomic regions affected by *CI* SVs revealed patterns that align with a north-south European cline and an east-west African cline, while no such associations were observed for *CII* SVs. We similarly observed that the recombination rates of *CI* SVs uniquely demonstrated Puerto Rican admixture. These results might indicate the potentially prominent role of regions affected by *CI* SVs in recent population divergence compared to those influenced by *CII* SVs. This further suggests a potential role for both homology and local sequence contexts in influencing recombination rates.

To support our investigation into repair and variation, particularly concerning human disease, we focused on mechanistic differences between common and rare SVs, including *de novo* SVs in healthy populations and rare disease cohorts. For this purpose, we utilized trios (parents and children) from the 1KG project, where we identified SVs that were common (AF >5%) and inherited, rare (AF <1%) and inherited, as well as *de novo* to a proband. Next, we compared cluster assignment frequencies and the underlying DSB repair mechanism patterns associated with germline *de novo* variants from individuals with rare diseases in the 100KG project(50). The participants included those with neurological and neurodevelopmental disorders (NND; n = 209), ultra-rare disorders (URD; n = 37), cardiovascular disorders (n = 24), renal and urinary tract disorders (RUT; n = 21), endocrine disorders (n = 20), and other disorders (n = 85). We posited that since *de novo* variants can significantly contribute to clinical phenotypes, the frequency of the repair mechanisms and the underlying features of SVs may differ between *de novo* SVs from individuals with rare diseases and both *de novo* and inherited SVs from healthy individuals.

Interestingly, we found an increasing proportion of common (31%), rare (44%), and *de novo* SVs (59%; **Figure 6A**) belonging to *CI* in the 1KG dataset. However, in most rare disease cohorts, the frequency of *CI* SVs was low, except for those with cardiovascular disorders (59%). Upon further examination of specific SVs, we discovered that the *CI_nHD_* variant, chr13-94711392-DEL-185, was present in multiple individuals, most of whom exhibited abnormal developmental phenotypes. Notably, this SV is located within SOX21, a transcription factor known to regulate neuronal differentiation(65); further, the SV perturbs pELS and CTCF sites (**Figure 6B**). However, we note that this SV was not identified as a causal variant for the diseases in these individuals. We then compared the underlying patterns of repair across these cohorts. Upon calculating the effect size between the cohorts using z-score normalized features, we found that when using the H1-hESC as a proxy, the upstream level of the histone marker H3K27me3 is distinct among individuals with URD (**Figure 6C**). We observed a pattern of decreasing effect sizes from inherited to *de novo* 1KG SVs. Since patients with URDs can display clinically complex phenotypes, we compared the reported phenotypes from the participants with *CI* and *CII de novo* SVs. We found that URD individuals with *CI* SVs displayed a non-significant enrichment in their abnormal genitourinary system and liver. In contrast, individuals with *CII* SVs have a non-significant enrichment in having abnormal skeletal and endocrine systems. We believe that larger sample sizes may further bolster clinical distinction between patients with URDs. This set of results highlights that while the repair mechanisms employed may not differ between healthy and diseased individuals, DNA accessibility associated features that contribute to effective repair may be altered, potentially reinforcing observed phenotypes.

**Figure 6.**
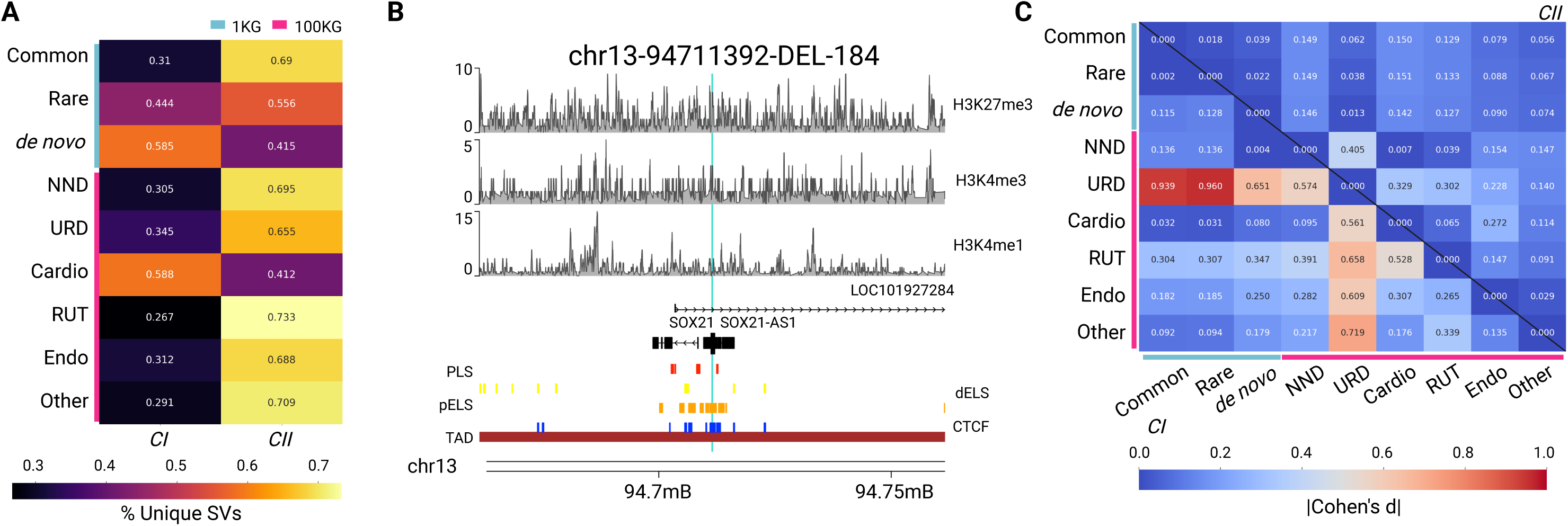
Application of clustering-based strategy to explore pathogenicity. **(A).** Heatmap showing *CI* and *CII* assignments across healthy (1KG) and diseased (100KG) cohorts, differentiated by turquoise and red bars. The healthy cohorts are represented by SVs from 1KG trios, including those that are commonly inherited, rare, or de novo. The diseased cohorts comprise *de novo* deletions from individuals with rare diseases, which include neurological and neurodevelopmental disorders (NND), ultra-rare disorders (URD), cardiovascular disorders (Cardio), renal and urinary tract disorders (RUT), endocrine disorders (Endo), and other disorders (n=85). Proportions are calculated row-wise. **(B).** Karyoplot of the SV, chr13-94711392-DEL-184, found in multiple individuals across the 100KG project cohorts. The chromosome’s ideogram is located at the bottom. Above are genomic elements, including the presence of a TAD, CTCF sites, pELS, dELS, and promoter-like signatures, along with the CDSs that the SV affects. The epigenetic context associated with this SV is represented above, as per H3K27me3, H3K4me3, and H3K4me1 in H1-hESCs. **(C).** Heatmap illustrating the magnitude of the effect sizes across healthy and diseased SV cohorts for the local upstream levels of H3K27me3. The upper and lower triangles display the *CII* and *CI* differences, respectively.

## Discussion

Recent advances in lrWGS and genome assembly algorithms have established accurate SV catalogs. Genomic consortia, including HGSVC(13), Human Pan Genome Reference Consortium (HPRC)(66), AllofUs(67), and 1KG-ONT(43) projects, are leveraging these technical advancements to describe SVs from diverse human populations. Beyond extending our understanding of how SVs influence genotype-phenotype relationships, these catalogs can be further utilized to investigate the mechanistic origins of SVs. In this work, we characterized the underlying repair mechanisms likely associated with the mechanistic origin of SVs in diverse human populations and some rare disease cohorts.

First, we applied a simple homology-based workflow to classify SVs into three categories (HLH, ILH, and NLH), inspired by DSB repair pathways (HR, SSA, aEJ, and NHEJ). We focused primarily on high-quality, de novo assembled haplotype-specific lrWGS SVs identified in the HGSVC2 study. Additionally, we utilized SVs from orthogonal datasets from lrWGS (1KG-ONT) and srWGS (high-coverage 1KG project) platforms to systematically identify patterns in SVs. Our observation that most SVs belong to the NLH class is consistent with previous reports that NHEJ occurs at the highest frequency, regardless of the cell cycle. However, we found that lrWGS can uncover more homology-driven SVs than srWGS due to its superior ability to detect SVs that span repetitive regions in the genome.

Subsequently, we quantified the propensity of SVs to affect distinct genomic elements and expectedly observed that most are under strong negative selection. However, we noted interesting deviations from this pattern for SVs affecting genes, specifically those belonging to certain pathways. For instance, we found that HLH/ILH and NLH SVs are significantly enriched for affecting genes involved in glycosylation, which include mucins, and the clearance of ligands by SR pathways, respectively. The relationship between homology-based SVs and mucin genes can be attributed to the high number of variable number tandem repeats present in these genes(56). In contrast, the enrichment of NLH SVs affecting genes involved in the clearance of ligands by SRs (i.e., lipoproteins and immunoglobulins) may indicate an association between gene function and repair mechanisms. It is well established that variation in immunoglobulins is introduced through V(D)J recombination, primarily conducted by NHEJ(57). However, these SVs are not expected in this germline context since V(D)J is a mechanism for introducing somatic variation. Therefore, this observation potentially points to a recognized technical artifact from sequencing lymphoblastoid cell lines (LCLs) characterized by a high rate of variants in IG loci(68). Regardless, future studies should investigate the association between repair choice, the introduction of SVs, and genes with high repeat element variability.

Furthermore, we observed significant differences in DNA shape, epigenomic, and sequence feature profiles among the simple homology-based SV classes. For example, we found that HLH SVs exhibited less variation but greater flanking MGW and more pronounced local methylation. This suggests that these regions are more homogeneous and rigid with high GC content. We therefore postulate that a region’s local context may influence DSB repair effectiveness and/or efficiency, and/or detection(69). Such contexts could be mediated by DNA accessibility and/or biophysical properties (for example, flanking MGW). These observations align with a prior study showing a higher DSB burden among Alzheimer’s patients, where genomic loci affected by SVs had greater DNA accessibility (70).

We also noted significant differences in additional features underlying SVs belonging to distinct homology-based categories. Therefore, we trained an active-learning-based unsupervised model to classify SVs, which considered homology, DNA shape, epigenetics, replication timing, and sequence profiles (including non-B DNA, GC content, and repeat annotations) local to or within the SV. To achieve this, we applied and optimized the HDBSCAN algorithm, demonstrating improved discrimination across these features while preserving homology patterns captured by two distinct sub-clusters *CI_HD_* (enriched among HLH SVs) and *CII_nHD_* (enriched among NLH SVs). Next, we characterized the ChromHMM states of SVs in distinct sub-clusters derived from our unsupervised approach. Interestingly, we found that *CI_nHD_* SVs are enriched in bivalently transcribed transcription start sites, which are relevant to neural development(71, 72). The importance of bivalent states and SV introduction may align with paradoxes aligned in previous studies(32, 33, 42), whereby it remains unclear whether homology-based repair mechanisms exclusively prefer to engage in repair in heterochromatin or euchromatin. This may also suggest that homology-based repair mechanisms, such as HR, SSA, or aEJ, preferentially target “operational” regions of the genome rather than quiescent regions.

Furthermore, we observed that deletions in the *CI_HD_* subcluster significantly disrupt local genomic folding compared to deletions in the *CII_nHD_* subcluster. These findings align with previous work showing that homology-based repair is enhanced when homology searching is limited by physical proximity(73–75). Proximity may play a role in repair kinetics since some DSB repair processes benefit from chromatin mobility(73); this can necessitate coordination between the repair machinery, the broken ends, and the nuclear periphery(76). Therefore, we posit that upon a repair attempt, DSB breakpoints for homology-driven deletions may start with greater physical proximity. However, following the introduction of a deletion, the flanking regions may require more significant relocation or reorganization. Further experimental work will be necessary to verify DNA interactions before and after HR and NHEJ to elucidate repair kinetics.

Next, we utilized rare deletions from the 1KG project to investigate genomic regions characterized by a high density of SVs associated with the *CI* or *CII*. For this analysis, we focused on genome-wide SV hotspots that are consistent across individuals from distinct ancestries. Interestingly, we discovered that *CI_HD_* SVs are enriched among genomic intervals primarily located in sub-telomeric regions. When considering sub-pathways of HR, sub-telomeres are often repaired using the error-prone Break-Induced Replication (BIR) pathway (64, 77), which is further relevant due to its clinical implications (69, 78), rather than the more accurate Synthesis-Dependent Strand Annealing (SDSA) pathway. Recent work has shown that pathogenic variants in the sub-telomeric region of chrX can be mediated through inverted repeats (which give rise to hairpins, a non-B DNA structure) and were likely introduced via BIR(69). Moreover, fork stalling and template switching/ microhomology-mediated BIR (FoSTeS/MMBIR) mechanism has been linked to rearrangements found on chromosome 17 among individuals with Potockli-Lupski microduplication syndrome (PTLS)(78). Similarly, there are several disorders with etiological links to faulty NAHR, for example, 15q13.3 microdeletion syndrome (22), Charcot-Marie-Tooth disease type 1A (79), X-linked ichthyosis (80), and 7q11.23 deletions in Williams-Beuren syndrome (81). Other relevant disorders, such as Fanconi anemia(20), which can cause defective HR, can present with phenotypes such as increased incidence of cancer, myelodysplasia, and congenital anomalies(82). Therefore, clinically, there is justification for probing SV introduction within the germline as a strategy to appreciate disease manifestation and refine variant interpretation. Although most of the germline variants we analyzed were from healthy individuals, we characterized repair attempts for only a subset of DSB sites present upon sequencing that meet our SV length threshold (50 bp). Therefore, since our study relies on identifiable repair records, we are limited to imperfect repair outcomes. It is reasonable to assume that *CI_HD_* SVs may be biased toward BIR instead of SDSA, given the location of the cluster’s hotspots we identified, and that SDSA is unlikely to leave behind evidence of its usage. To mitigate this selection bias, higher throughput and more specific reporter assays can be applied to confirm the underlying features of repair in both perfect and imperfect repair scenarios, thereby capturing the entire spectrum of homology-based repair pathways and sub-pathways.

Finally, we applied our unsupervised method to classify common, rare, and *de novo* variants in healthy individuals (1KG project) and those with rare diseases (100KG study). Most *de novo* SVs in rare disease cohorts exhibited a higher frequency of *CII* SVs. However, we noticed that *de novo CI* and *CII* SVs in individuals with URDs could be distinguished based on their local levels of H3K27me3 signals compared to SVs from other rare diseases and healthy individuals. This observation is intriguing, considering that URDs are primarily monogenic and exhibit complex clinical presentations. This suggests that while SVs from patients with URDs may be epigenetically distinct, this finding likely does not play a critical role in disease etiology, but it remains noteworthy. With more samples, the relationship between DSB repair mechanisms and germline SVs could aid in classifying clinical phenotypes in rare diseases.

Overall, our study comprehensively characterized DSB repair mechanisms and their associated features underlying the mechanistic origins of SVs. In contrast to previous studies that primarily relied on large deletions from srWGS, our work utilized multiple population-level SV repositories from various lrWGS and srWGS efforts. Additionally, we considered various epigenomic, DNA shape, and sequence-related characteristics to gain insights into distinct classes of SV mechanisms. This study builds upon earlier computational efforts, which predominantly utilized sequence homology and repeat annotations. Despite several significant findings, we acknowledge certain technical limitations in our study. For instance, we obtained 1975 bp upstream and downstream sequences based on the reference genome when calculating local sequence homology. While effective, this approach overlooks potential haplotype-specific variation. Consequently, locally assembled and consensus sequences could improve the accuracy of these alignments. Furthermore, our analyses concentrated solely on insertions and deletions. We note that we characterized inversions within the HGSVC2 and 1KG-ONT datasets. Although, given the small sample sizes and homogeneity in their threshold-based label (ie. almost all were labelled as NLH), we could not draw comparisons across SV classes and deploy an unsupervised model. Future computational work should broaden the range of SV types to include complex SVs. Additionally, we employed *de novo* SVs from the 1 KG project to characterize and compare the usage of DSB repair mechanisms in healthy individuals and those with rare diseases. However, it is presumed that the rate of *de novo* variants from the 1 KG project is unexpectedly high (44), at approximately 10 times the expected rate, with approximately 100 de novo mutations per child. This indicates that the origin of the *de novo* SVs in this dataset reflects another technical artifact from the LCLs used during sequencing, rather than an accurate representation of mechanisms deployed in the germline or during development. Finally, we used H1-hESCs as a proxy for epigenetic and replication timing annotations of SVs(53). This assumes that the chromatin profiles of H1 cells are well conserved and exhibit minimal dynamics during the acquisition of SVs. To determine if patterns of DNA accessibility and repair choice have a correlational or causative relationship, future studies should enhance this approach by employing multi-omics data to investigate the epigenomic states of regions proximal to SVs in healthy individuals and those with rare diseases. For example, this could include characterizing 3D genome folding, histone modifications, and DNA accessibility proximal to variants in affected and unaffected siblings using Hi-C and ATAC-Seq assays.

Despite the outlined limitations, our study presents the most thorough computational characterization of DSB repair mechanisms. Our study identifies underlying features that influence the mechanistic origin of SVs in diverse human populations. We anticipate that continued efforts to investigate features that support (or hinder) the introduction of SVs will help us accurately delineate their role in defining genotype-phenotype relationships.

## Source code and data availability

The HGSVC2 and high-coverage 1000 genome phase 3 SV datasets were downloaded from the IGSR data portal (https://ftp.1000genomes.ebi.ac.uk/vol1/ftp/data_collections/HGSVC2/release/v2.0/integrated_c allset/ & http://ftp.1000genomes.ebi.ac.uk/vol1/ftp/data_collections/1000G_2504_high_coverage/working/20220422_3202_phased_SNV_INDEL_SV/).

Additionally, epigenomics and cCRE annotations (https://screen.encodeproject.org/) are available through the ENCODE project data portal. 1KG-ONT SVs are hosted on Amazon bucket (https://s3.amazonaws.com/1000g-ont/Gustafson_etal_2024_preprint_SUPPLEMENTAL/20240423_jasmine_intrasample_noBND_custom_suppvec_alphanumeric_header_JASMINE.vcf.gz). The 100KG project data utilized in this study can be accessed by logging into the Genomics England research environment.

Finally, relevant source codes for this study, including scripts for simulation, annotation, model training, and downstream analyses, can be accessed on the project GitHub page(https://github.com/kumarlab-compomics/SV_mechanism).

## Acknowledgments

This research was made possible through access to data in the National Genomic Research Library, which is managed by Genomics England Limited (a wholly owned company of the Department of Health and Social Care). The National Genomic Research Library holds data provided by patients and collected by the NHS as part of their care and data collected as part of their participation in research. The National Genomic Research Library is funded by the National Institute for Health Research and NHS England. The Wellcome Trust, Cancer Research UK, and the Medical Research Council have also funded research infrastructure.

## References

1. Kumar, S. and Gerstein, M. (2023) Unified views on variant impact across many diseases. Trends Genet., 39, 442–450.

2. 100, 000 Genomes Project Pilot Investigators, Smedley, D., Smith, K.R., Martin, A., Thomas, E.A., McDonagh, E.M., Cipriani, V., Ellingford, J.M., Arno, G., Tucci, A., et al. (2021) 100, 000 genomes pilot on rare-disease diagnosis in health care - preliminary report. N. Engl. J. Med., 385, 1868–1880.

3. Pang, A.W., MacDonald, J.R., Pinto, D., Wei, J., Rafiq, M.A., Conrad, D.F., Park, H., Hurles, M.E., Lee, C., Venter, J.C., et al. (2010) Towards a comprehensive structural variation map of an individual human genome. Genome Biol., 11, R52.

4. Scott, A.J., Chiang, C. and Hall, I.M. (2021) Structural variants are a major source of gene expression differences in humans and often affect multiple nearby genes. Genome Res., 31, 2249–2257.

5. Abel, H.J., Larson, D.E., Regier, A.A., Chiang, C., Das, I., Kanchi, K.L., Layer, R.M., Neale, B.M., Salerno, W.J., Reeves, C., et al. (2020) Mapping and characterization of structural variation in 17, 795 human genomes. Nature, 583, 83–89.

6. Carvalho, C.M.B. and Lupski, J.R. (2016) Mechanisms underlying structural variant formation in genomic disorders. Nat. Rev. Genet., 17, 224–238.

7. Merker, J.D., Wenger, A.M., Sneddon, T., Grove, M., Zappala, Z., Fresard, L., Waggott, D., Utiramerur, S., Hou, Y., Smith, K.S., et al. (2018) Long-read genome sequencing identifies causal structural variation in a Mendelian disease. Genet. Med., 20, 159–163.

8. Trost, B., Thiruvahindrapuram, B., Chan, A.J.S., Engchuan, W., Higginbotham, E.J., Howe, J.L., Loureiro, L.O., Reuter, M.S., Roshandel, D., Whitney, J., et al. (2022) Genomic architecture of autism from comprehensive whole-genome sequence annotation. Cell, 185, 4409–4427.e18.

9. Kosugi, S. and Terao, C. (2024) Comparative evaluation of SNVs, indels, and structural variations detected with short- and long-read sequencing data. Hum. Genome Var., 11, 18.

10. Mahmoud, M., Gobet, N., Cruz-Dávalos, D.I., Mounier, N., Dessimoz, C. and Sedlazeck, F.J. (2019) Structural variant calling: the long and the short of it. Genome Biol., 20, 246.

11. Jeong, H., Dishuck, P.C., Yoo, D., Harvey, W.T., Munson, K.M., Lewis, A.P., Kordosky, J., Garcia, G.H., Human Genome Structural Variation Consortium (HGSVC), Yilmaz, F., et al. (2025) Structural polymorphism and diversity of human segmental duplications. Nat. Genet., 57, 390–401.

12. Bolognini, D., Halgren, A., Lou, R.N., Raveane, A., Rocha, J.L., Guarracino, A., Soranzo, N., Chin, C.-S., Garrison, E. and Sudmant, P.H. (2024) Recurrent evolution and selection shape structural diversity at the amylase locus. Nature, 634, 617–625.

13. Ebert, P., Audano, P.A., Zhu, Q., Rodriguez-Martin, B., Porubsky, D., Bonder, M.J., Sulovari, A., Ebler, J., Zhou, W., Serra Mari, R., et al. (2021) Haplotype-resolved diverse human genomes and integrated analysis of structural variation. Science, 372.

14. Miller, D.E., Sulovari, A., Wang, T., Loucks, H., Hoekzema, K., Munson, K.M., Lewis, A.P., Fuerte, E.P.A., Paschal, C.R., Walsh, T., et al. (2021) Targeted long-read sequencing identifies missing disease-causing variation. Am. J. Hum. Genet., 108, 1436–1449.

15. Kieffer, S.R. and Lowndes, N.F. (2022) Immediate-Early, Early, and Late responses to DNA double stranded breaks. Front. Genet., 13, 793884.

16. Ballinger, T.J., Bouwman, B.A.M., Mirzazadeh, R., Garnerone, S., Crosetto, N. and Semple, C.A. (2019) Modeling double strand break susceptibility to interrogate structural variation in cancer. Genome Biol., 20, 28.

17. Dileep, V., Boix, C.A., Mathys, H., Marco, A., Welch, G.M., Meharena, H.S., Loon, A., Jeloka, R., Peng, Z., Bennett, D.A., et al. (2023) Neuronal DNA double-strand breaks lead to genome structural variations and 3D genome disruption in neurodegeneration. Cell, 186, 4404–4421.e20.

18. Aguilera, A. and García-Muse, T. (2013) Causes of genome instability. Annu. Rev. Genet., 47, 1–32.

19. O’Driscoll, M., Gennery, A.R., Seidel, J., Concannon, P. and Jeggo, P.A. (2004) An overview of three new disorders associated with genetic instability: LIG4 syndrome, RS-SCID and ATR-Seckel syndrome. DNA Repair (Amst.), 3, 1227–1235.

20. Peake, J.D. and Noguchi, E. (2022) Fanconi anemia: current insights regarding epidemiology, cancer, and DNA repair. Hum. Genet., 141, 1811–1836.

21. Sharma, R., Lewis, S. and Wlodarski, M.W. (2020) DNA Repair Syndromes and Cancer: Insights Into Genetics and Phenotype Patterns. Front Pediatr, 8, 570084.

22. Sharp, A.J., Mefford, H.C., Li, K., Baker, C., Skinner, C., Stevenson, R.E., Schroer, R.J., Novara, F., De Gregori, M., Ciccone, R., et al. (2008) A recurrent 15q13.3 microdeletion syndrome associated with mental retardation and seizures. Nat. Genet., 40, 322–328.

23. Farmer, H., McCabe, N., Lord, C.J., Tutt, A.N.J., Johnson, D.A., Richardson, T.B., Santarosa, M., Dillon, K.J., Hickson, I., Knights, C., et al. (2005) Targeting the DNA repair defect in BRCA mutant cells as a therapeutic strategy. Nature, 434, 917–921.

24. Pilié, P.G., Gay, C.M., Byers, L.A., O’Connor, M.J. and Yap, T.A. (2019) PARP inhibitors: Extending benefit beyond BRCA-mutant cancers. Clin. Cancer Res., 25, 3759–3771.

25. Abyzov, A., Li, S., Kim, D.R., Mohiyuddin, M., Stütz, A.M., Parrish, N.F., Mu, X.J., Clark, W., Chen, K., Hurles, M., et al. (2015) Analysis of deletion breakpoints from 1, 092 humans reveals details of mutation mechanisms. Nat. Commun., 6, 7256.

26. Lam, H.Y.K., Mu, X.J., Stütz, A.M., Tanzer, A., Cayting, P.D., Snyder, M., Kim, P.M., Korbel, J.O. and Gerstein, M.B. (2010) Nucleotide-resolution analysis of structural variants using BreakSeq and a breakpoint library. Nat. Biotechnol., 28, 47–55.

27. Pang, A.W.C., Migita, O., Macdonald, J.R., Feuk, L. and Scherer, S.W. (2013) Mechanisms of formation of structural variation in a fully sequenced human genome. Hum. Mutat., 34, 345–354.

28. Ortega, P., Gómez-González, B. and Aguilera, A. (2021) Heterogeneity of DNA damage incidence and repair in different chromatin contexts. DNA Repair, 107, 103210.

29. Caron, P., Pobega, E. and Polo, S.E. (2021) DNA Double-Strand Break Repair: All Roads Lead to HeterochROMAtin Marks. Front. Genet., 12, 730696.

30. Crosetto, N., Mitra, A., Silva, M.J., Bienko, M., Dojer, N., Wang, Q., Karaca, E., Chiarle, R., Skrzypczak, M., Ginalski, K., et al. (2013) Nucleotide-resolution DNA double-strand break mapping by next-generation sequencing. Nat. Methods, 10, 361–365.

31. Akhtar, W., de Jong, J., Pindyurin, A.V., Pagie, L., Meuleman, W., de Ridder, J., Berns, A., Wessels, L.F.A., van Lohuizen, M. and van Steensel, B. (2013) Chromatin position effects assayed by thousands of reporters integrated in parallel. Cell, 154, 914–927.

32. Schep, R., Brinkman, E.K., Leemans, C., Vergara, X., van der Weide, R.H., Morris, B., van Schaik, T., Manzo, S.G., Peric-Hupkes, D., van den Berg, J., et al. (2021) Impact of chromatin context on Cas9-induced DNA double-strand break repair pathway balance. Mol. Cell, 81, 2216–2230.e10.

33. Clouaire, T., Rocher, V., Lashgari, A., Arnould, C., Aguirrebengoa, M., Biernacka, A., Skrzypczak, M., Aymard, F., Fongang, B., Dojer, N., et al. (2018) Comprehensive mapping of histone modifications at DNA double-strand breaks deciphers repair pathway chromatin signatures. Mol. Cell, 72, 250–262.e6.

34. Vítor, A.C., Huertas, P., Legube, G. and de Almeida, S.F. (2020) Studying DNA double-strand break repair: An ever-growing toolbox. Front. Mol. Biosci., 7, 24.

35. Liu, S.-C., Feng, Y.-L., Sun, X.-N., Chen, R.-D., Liu, Q., Xiao, J.-J., Zhang, J.-N., Huang, Z.-C., Xiang, J.-F., Chen, G.-Q., et al. (2022) Target residence of Cas9-sgRNA influences DNA double-strand break repair pathway choices in CRISPR/Cas9 genome editing. Genome Biol., 23, 165.

36. Pallaseni, A., Peets, E.M., Girling, G., Crepaldi, L., Kuzmin, I., Moor, M., Muñoz-Subirana, N., Schimmel, J., Serçin, Ö., Mardin, B.R., et al. (2024) The interplay of DNA repair context with target sequence predictably biases Cas9-generated mutations. Nat. Commun., 15, 10271.

37. Hufnagl, A., Herr, L., Friedrich, T., Durante, M., Taucher-Scholz, G. and Scholz, M. (2015) The link between cell-cycle dependent radiosensitivity and repair pathways: a model based on the local, sister-chromatid conformation dependent switch between NHEJ and HR. DNA Repair (Amst.), 27, 28–39.

38. Fortini, P., Ferretti, C. and Dogliotti, E. (2013) The response to DNA damage during differentiation: pathways and consequences. Mutat. Res., **743–744**, 160–168.

39. Beerman, I., Seita, J., Inlay, M.A., Weissman, I.L. and Rossi, D.J. (2014) Quiescent hematopoietic stem cells accumulate DNA damage during aging that is repaired upon entry into cell cycle. Cell Stem Cell, 15, 37–50.

40. Gnügge, R., Reginato, G., Cejka, P. and Symington, L.S. (2023) Sequence and chromatin features guide DNA double-strand break resection initiation. Mol. Cell, 83, 1237–1250.e15.

41. Yin, X., Liu, M., Tian, Y., Wang, J. and Xu, Y. (2017) Cryo-EM structure of human DNA-PK holoenzyme. Cell Res., 27, 1341–1350.

42. Kendek, A., Wensveen, M.R. and Janssen, A. (2021) The sound of silence: How silenced chromatin orchestrates the repair of double-strand breaks. Genes (Basel), 12, 1415.

43. Gustafson, J.A., Gibson, S.B., Damaraju, N., Zalusky, M.P.G., Hoekzema, K., Twesigomwe, D., Yang, L., Snead, A.A., Richmond, P.A., De Coster, W., et al. (2024) High-coverage nanopore sequencing of samples from the 1000 Genomes Project to build a comprehensive catalog of human genetic variation. Genome Res., 34, 2061–2073.

44. Byrska-Bishop, M., Evani, U.S., Zhao, X., Basile, A.O., Abel, H.J., Regier, A.A., Corvelo, A., Clarke, W.E., Musunuri, R., Nagulapalli, K., et al. (2022) High-coverage whole-genome sequencing of the expanded 1000 Genomes Project cohort including 602 trios. Cell, 185, 3426–3440.e19.

45. Dumont, A., Mendiboure, N., Savocco, J., Anani, L., Moreau, P., Thierry, A., Modolo, L., Jost, D. and Piazza, A. (2024) Mechanism of homology search expansion during recombinational DNA break repair in Saccharomyces cerevisiae. Mol. Cell, 84, 3237–3253.e6.

46. Aleksandrov, R., Hristova, R., Stoynov, S. and Gospodinov, A. (2020) The Chromatin Response to Double-Strand DNA Breaks and Their Repair. Cells, 9.

47. Setton, J., Hadi, K., Choo, Z.-N., Kuchin, K.S., Tian, H., Da Cruz Paula, A., Rosiene, J., Selenica, P., Behr, J., Yao, X., et al. (2023) Long-molecule scars of backup DNA repair in BRCA1- and BRCA2-deficient cancers. Nature, 621, 129–137.

48. Blasiak, J. (2021) Single-strand annealing in cancer. Int. J. Mol. Sci., 22, 2167.

49. Ceccaldi, R., Rondinelli, B. and D’Andrea, A.D. (2016) Repair pathway choices and consequences at the double-strand break. Trends Cell Biol., 26, 52–64.

50. The National Genomics Research Library v5.1, Genomics England Genomics England, 1.

51. Lee, S.-Y., Wang, H., Cho, H.J., Xi, R. and Kim, T.-M. (2022) The shaping of cancer genomes with the regional impact of mutation processes. Exp. Mol. Med., 54, 1049–1060.

52. Roadmap Epigenomics Consortium, Kundaje, A., Meuleman, W., Ernst, J., Bilenky, M., Yen, A., Heravi-Moussavi, A., Kheradpour, P., Zhang, Z., Wang, J., et al. (2015) Integrative analysis of 111 reference human epigenomes. Nature, 518, 317–330.

53. Thomson, J.A., Itskovitz-Eldor, J., Shapiro, S.S., Waknitz, M.A., Swiergiel, J.J., Marshall, V.S. and Jones, J.M. (1998) Embryonic stem cell lines derived from human blastocysts. Science, 282, 1145–1147.

54. Jeffares, D.C., Jolly, C., Hoti, M., Speed, D., Shaw, L., Rallis, C., Balloux, F., Dessimoz, C., Bähler, J. and Sedlazeck, F.J. (2017) Transient structural variations have strong effects on quantitative traits and reproductive isolation in fission yeast. Nat. Commun., 8, 14061.

55. Collins, R.L., Brand, H., Karczewski, K.J., Zhao, X., Alföldi, J., Francioli, L.C., Khera, A.V., Lowther, C., Gauthier, L.D., Wang, H., et al. (2020) A structural variation reference for medical and population genetics. Nature, 581, 444–451.

56. Fowler, J., Vinall, L. and Swallow, D. (2001) Polymorphism of the human muc genes. Front. Biosci., 6, D1207–15.

57. Nicolas, L., Cols, M., Choi, J.E., Chaudhuri, J. and Vuong, B. (2018) Generating and repairing genetically programmed DNA breaks during immunoglobulin class switch recombination. F1000Res., 7, 458.

58. Lachenbruch, P.A. and Cohen, J. (1989) Statistical Power Analysis for the Behavioral Sciences (2nd ed.). J. Am. Stat. Assoc., 84, 1096.

59. Campello, R.J.G.B., Moulavi, D. and Sander, J. (2013) Density-based clustering based on hierarchical density estimates. In *Advances in Knowledge Discovery and Data Mining*, Lecture notes in computer science. Springer Berlin Heidelberg, Berlin, Heidelberg, pp. 160–172.

60. Liu, Y., Sung, S., Kim, Y., Li, F., Gwon, G., Jo, A., Kim, A.-K., Kim, T., Song, O.-K., Lee, S.E., et al. (2016) ATP-dependent DNA binding, unwinding, and resection by the Mre11/Rad50 complex. EMBO J., 35, 743–758.

61. Zhou, J. (2022) Sequence-based modeling of three-dimensional genome architecture from kilobase to chromosome scale. Nat. Genet., 54, 725–734.

62. Gunsalus, L.M., McArthur, E., Gjoni, K., Kuang, S., Pittman, M., Capra, J.A. and Pollard, K.S. (2023) Comparing chromatin contact maps at scale: methods and insights. bioRxivorg, 10.1101/2023.04.04.535480.

63. Spence, J.P. and Song, Y.S. (2019) Inference and analysis of population-specific fine-scale recombination maps across 26 diverse human populations. Sci. Adv., 5, eaaw9206.

64. Batté, A., Brocas, C., Bordelet, H., Hocher, A., Ruault, M., Adjiri, A., Taddei, A. and Dubrana, K. (2017) Recombination at subtelomeres is regulated by physical distance, double-strand break resection and chromatin status. EMBO J., 36, 2609–2625.

65. Whittington, N., Cunningham, D., Le, T.-K., De Maria, D. and Silva, E.M. (2015) Sox21 regulates the progression of neuronal differentiation in a dose-dependent manner. Dev. Biol., 397, 237–247.

66. Liao, W.-W., Asri, M., Ebler, J., Doerr, D., Haukness, M., Hickey, G., Lu, S., Lucas, J.K., Monlong, J., Abel, H.J., et al. (2023) A draft human pangenome reference. Nature, 617, 312–324.

67. All of Us Research Program Genomics Investigators (2024) Genomic data in the All of Us Research Program. Nature, 627, 340–346.

68. Rodriguez, O.L., Gibson, W.S., Parks, T., Emery, M., Powell, J., Strahl, M., Deikus, G., Auckland, K., Eichler, E.E., Marasco, W.A., et al. (2020) A novel framework for characterizing genomic haplotype diversity in the human immunoglobulin heavy chain locus. Front. Immunol., 11, 2136.

69. Grochowski, C.M., Bengtsson, J.D., Du, H., Gandhi, M., Lun, M.Y., Mehaffey, M.G., Park, K., Höps, W., Benito, E., Hasenfeld, P., et al. (2024) Inverted triplications formed by iterative template switches generate structural variant diversity at genomic disorder loci. Cell Genom., 4, 100590.

70. Zhang, X., Liu, Y., Huang, M., Gunewardena, S., Haeri, M., Swerdlow, R.H. and Wang, N. (2023) Landscape of double-stranded DNA breaks in postmortem brains from Alzheimer’s disease and non-demented individuals. J. Alzheimers. Dis., 94, 519–535.

71. Ramesh, V., Liu, F., Minto, M.S., Chan, U. and West, A.E. (2023) Bidirectional regulation of postmitotic H3K27me3 distributions underlie cerebellar granule neuron maturation dynamics. Elife, 12.

72. Liu, J., Wu, X., Zhang, H., Pfeifer, G.P. and Lu, Q. (2017) Dynamics of RNA polymerase II pausing and bivalent histone H3 methylation during neuronal differentiation in brain development. Cell Rep., 20, 1307–1318.

73. García Fernández, F., Almayrac, E., Carré Simon, À., Batrin, R., Khalil, Y., Boissac, M. and Fabre, E. (2022) Global chromatin mobility induced by a DSB is dictated by chromosomal conformation and defines the HR outcome. Elife, 11.

74. Lee, C.-S., Wang, R.W., Chang, H.-H., Capurso, D., Segal, M.R. and Haber, J.E. (2016) Chromosome position determines the success of double-strand break repair. Proc. Natl. Acad. Sci. U. S. A., 113, E146–54.

75. Roychowdhury, T. and Abyzov, A. (2019) Chromatin organization modulates the origin of heritable structural variations in human genome. Nucleic Acids Res., 47, 2766–2777.

76. Shokrollahi, M., Stanic, M., Hundal, A., Chan, J.N.Y., Urman, D., Jordan, C.A., Hakem, A., Espin, R., Hao, J., Krishnan, R., et al. (2024) DNA double-strand break-capturing nuclear envelope tubules drive DNA repair. Nat. Struct. Mol. Biol., 31, 1319–1330.

77. Chung, D.K.C., Chan, J.N.Y., Strecker, J., Zhang, W., Ebrahimi-Ardebili, S., Lu, T., Abraham, K.J., Durocher, D. and Mekhail, K. (2015) Perinuclear tethers license telomeric DSBs for a broad kinesin- and NPC-dependent DNA repair process. Nat. Commun., 6, 7742.

78. Zhang, F., Khajavi, M., Connolly, A.M., Towne, C.F., Batish, S.D. and Lupski, J.R. (2009) The DNA replication FoSTeS/MMBIR mechanism can generate genomic, genic and exonic complex rearrangements in humans. Nat. Genet., 41, 849–853.

79. Pentao, L., Wise, C.A., Chinault, A.C., Patel, P.I. and Lupski, J.R. (1992) Charcot-Marie-Tooth type 1A duplication appears to arise from recombination at repeat sequences flanking the 1.5 Mb monomer unit. Nat. Genet., 2, 292–300.

80. Van Esch, H., Hollanders, K., Badisco, L., Melotte, C., Van Hummelen, P., Vermeesch, J.R., Devriendt, K., Fryns, J.-P., Marynen, P. and Froyen, G. (2005) Deletion of VCX-A due to NAHR plays a major role in the occurrence of mental retardation in patients with X-linked ichthyosis. Hum. Mol. Genet., 14, 1795–1803.

81. Urbán, Z., Helms, C., Fekete, G., Csiszár, K., Bonnet, D., Munnich, A., Donis-Keller, H. and Boyd, C.D. (1996) 7q11.23 deletions in Williams syndrome arise as a consequence of unequal meiotic crossover. Am. J. Hum. Genet., 59, 958–962.

82. Glanz, A. and Fraser, F.C. (1982) Spectrum of anomalies in Fanconi anaemia. J. Med. Genet., 19, 412–416.

83. Kirsche, M., Prabhu, G., Sherman, R., Ni, B., Battle, A., Aganezov, S. and Schatz, M.C. (2023) Jasmine and Iris: population-scale structural variant comparison and analysis. Nat. Methods, 10.1038/s41592-022-01753-3.

84. Poplin, R., Ruano-Rubio, V., DePristo, M.A., Fennell, T.J., Carneiro, M.O., Van der Auwera, G.A., Kling, D.E., Gauthier, L.D., Levy-Moonshine, A., Roazen, D., et al. (2017) Scaling accurate genetic variant discovery to tens of thousands of samples. bioRxiv, 10.1101/201178.

85. Larson, D.E., Abel, H.J., Chiang, C., Badve, A., Das, I., Eldred, J.M., Layer, R.M. and Hall, I.M. (2019) Svtools: Population-scale analysis of structural variation. Bioinformatics, 35, 4782– 4787.

86. Chen, X., Schulz-Trieglaff, O., Shaw, R., Barnes, B., Schlesinger, F., Källberg, M., Cox, A.J., Kruglyak, S. and Saunders, C.T. (2016) Manta: rapid detection of structural variants and indels for germline and cancer sequencing applications. Bioinformatics, 32, 1220–1222.

87. Alexandrov, L.B., Nik-Zainal, S., Wedge, D.C., Campbell, P.J. and Stratton, M.R. (2013) Deciphering signatures of mutational processes operative in human cancer. Cell Rep., 3, 246–259.

88. Sloan, C.A., Chan, E.T., Davidson, J.M., Malladi, V.S., Strattan, J.S., Hitz, B.C., Gabdank, I., Narayanan, A.K., Ho, M., Lee, B.T., et al. (2016) ENCODE data at the ENCODE portal. Nucleic Acids Res., 44, D726–32.

89. Zhao, P.A., Sasaki, T. and Gilbert, D.M. (2020) High-resolution Repli-Seq defines the temporal choreography of initiation, elongation and termination of replication in mammalian cells. Genome Biol., 21, 76.

90. Reiff, S.B., Schroeder, A.J., Kırlı, K., Cosolo, A., Bakker, C., Mercado, L., Lee, S., Veit, A.D., Balashov, A.K., Vitzthum, C., et al. (2022) The 4D Nucleome Data Portal as a resource for searching and visualizing curated nucleomics data. Nat. Commun., 13, 2365.

91. Brison, O., El-Hilali, S., Azar, D., Koundrioukoff, S., Schmidt, M., Nähse, V., Jaszczyszyn, Y., Lachages, A.-M., Dutrillaux, B., Thermes, C., et al. (2019) Transcription-mediated organization of the replication initiation program across large genes sets common fragile sites genome-wide. Nat. Commun., 10, 5693.

92. Cer, R.Z., Donohue, D.E., Mudunuri, U.S., Temiz, N.A., Loss, M.A., Starner, N.J., Halusa, G.N., Volfovsky, N., Yi, M., Luke, B.T., et al. (2013) Non-B DB v2.0: a database of predicted non-B DNA-forming motifs and its associated tools. Nucleic Acids Res., 41, D94–D100.

93. Chiu, T.-P., Comoglio, F., Zhou, T., Yang, L., Paro, R. and Rohs, R. (2016) DNAshapeR: an R/Bioconductor package for DNA shape prediction and feature encoding. Bioinformatics, 32, 1211–1213.

94. Quinlan, A.R. and Hall, I.M. (2010) BEDTools: a flexible suite of utilities for comparing genomic features. Bioinformatics, 26, 841–842.

95. ENCODE Project Consortium, Moore, J.E., Purcaro, M.J., Pratt, H.E., Epstein, C.B., Shoresh, N., Adrian, J., Kawli, T., Davis, C.A., Dobin, A., et al. (2020) Expanded encyclopaedias of DNA elements in the human and mouse genomes. Nature, 583, 699–710.

96. Dixon, J.R., Jung, I., Selvaraj, S., Shen, Y., Antosiewicz-Bourget, J.E., Lee, A.Y., Ye, Z., Kim, A., Rajagopal, N., Xie, W., et al. (2015) Chromatin architecture reorganization during stem cell differentiation. Nature, 518, 331–336.

97. Shah, P.P., Keough, K.C., Gjoni, K., Santini, G.T., Abdill, R.J., Wickramasinghe, N.M., Dundes, C.E., Karnay, A., Chen, A., Salomon, R.E.A., et al. (2023) An atlas of lamina-associated chromatin across twelve human cell types reveals an intermediate chromatin subtype. Genome Biol., 24, 16.

98. Kumar, R., Nagpal, G., Kumar, V., Usmani, S.S., Agrawal, P. and Raghava, G.P.S. (2019) HumCFS: a database of fragile sites in human chromosomes. BMC Genomics, 19, 985.

99. Milacic, M., Beavers, D., Conley, P., Gong, C., Gillespie, M., Griss, J., Haw, R., Jassal, B., Matthews, L., May, B., et al. (2024) The reactome pathway knowledgebase 2024. Nucleic Acids Res., 52, D672–D678.

