## Supplementary material for "DNA shape and epigenomics distinguish the mechanistic origin of human genomic structural variations": SI Figures

**Table 1. Standardized residuals amongst homology-based labels and ChromHMM states.** Comparison of $\chi^2$- test results amongst states represented in all three workflow classes, for deletions (A) and insertions (B).

**Table 2. Description of z-score-derived features calculated using real and matched simulations SVs.**

**Table 3. Results from preprocessing and hyperparameter selection as per HDBSCAN optimization efforts.** Reflects evaluation metrics from valid clusters derived via different combinations in the number of PCA dimensions, minimum number of SVs per cluster, and distance metric. Reported results include the proportion of high-quality SVs amongst homology based classes, silhouette scores and Odds Ratio when comparing pseudo-HLH and pseudo-NLH clusters, for the deletions (A) and insertion (B) models.

**Table 4. Z-score-derived features included in the optimal HDBSCAN deletions (A) and insertions (B) models.**

**Table 5. Standardized residuals amongst cluster-based labels from deletions (A) and insertions (B) among ChromHMM states.** Comparison of $\chi^2$- test results amongst states represented in subcluster- based classes.

**Table 6. Summary table of main findings**

**Table 7. Glossary of full and abbreviated terms**

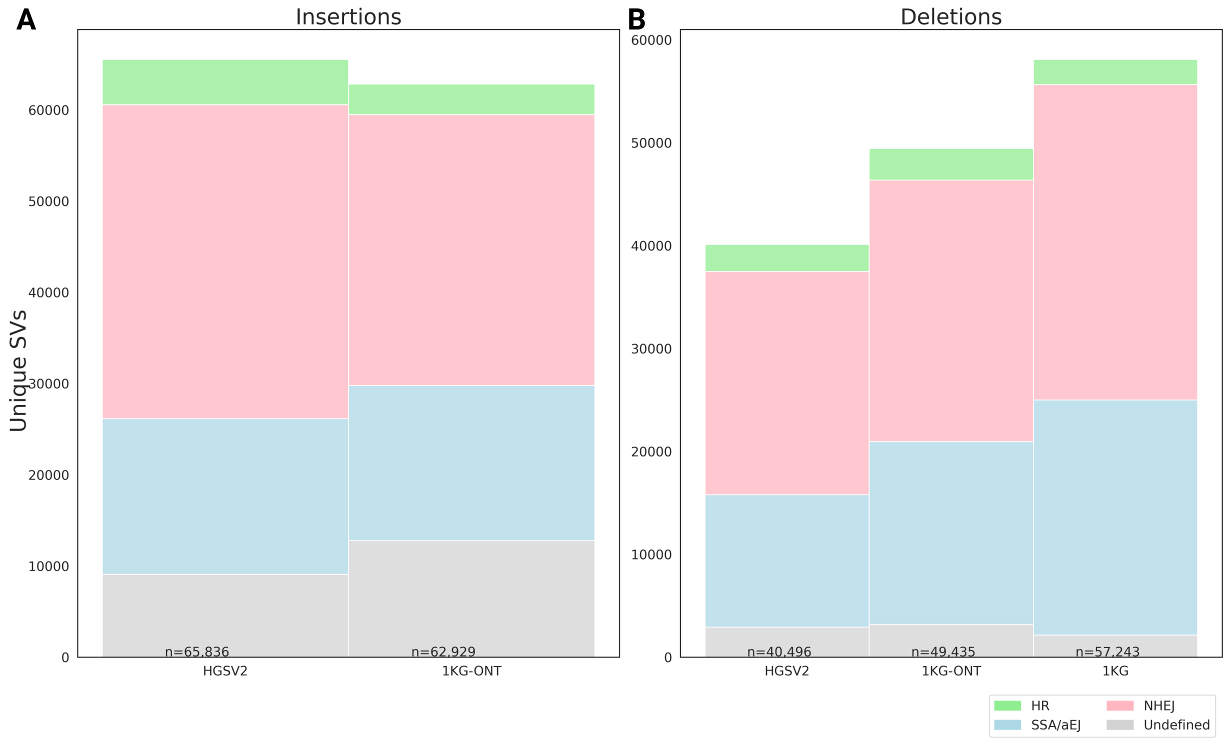


**Figure S1. Barplot depicting the absolute counts of insertions (A) and deletions (B) amongst the unique SVs in lrWGS (HGSVC2 & 1KG-ONT) and srWGS (high coverage 1KG phase 3) datasets.**

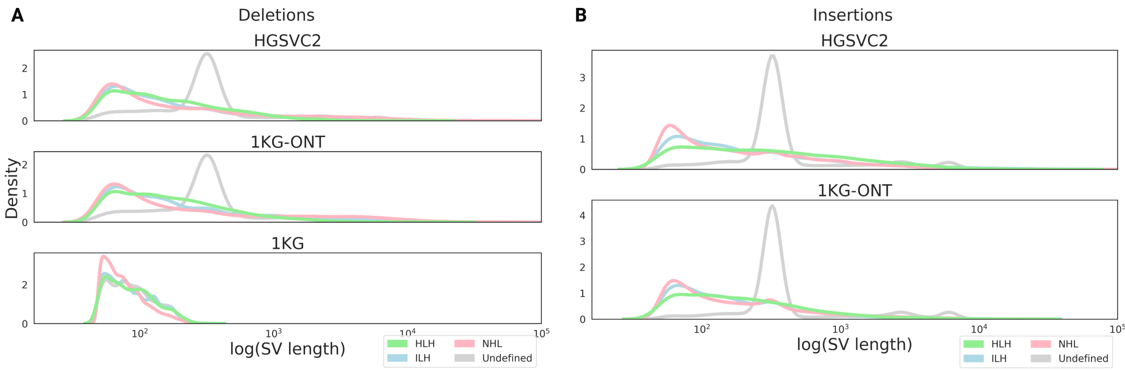


**Figure S2. Distribution of SV lengths by workflow-based label, across the datasets and SV types, deletions (A) and insertions (B).** Lengths depicted in logarithmic scale.

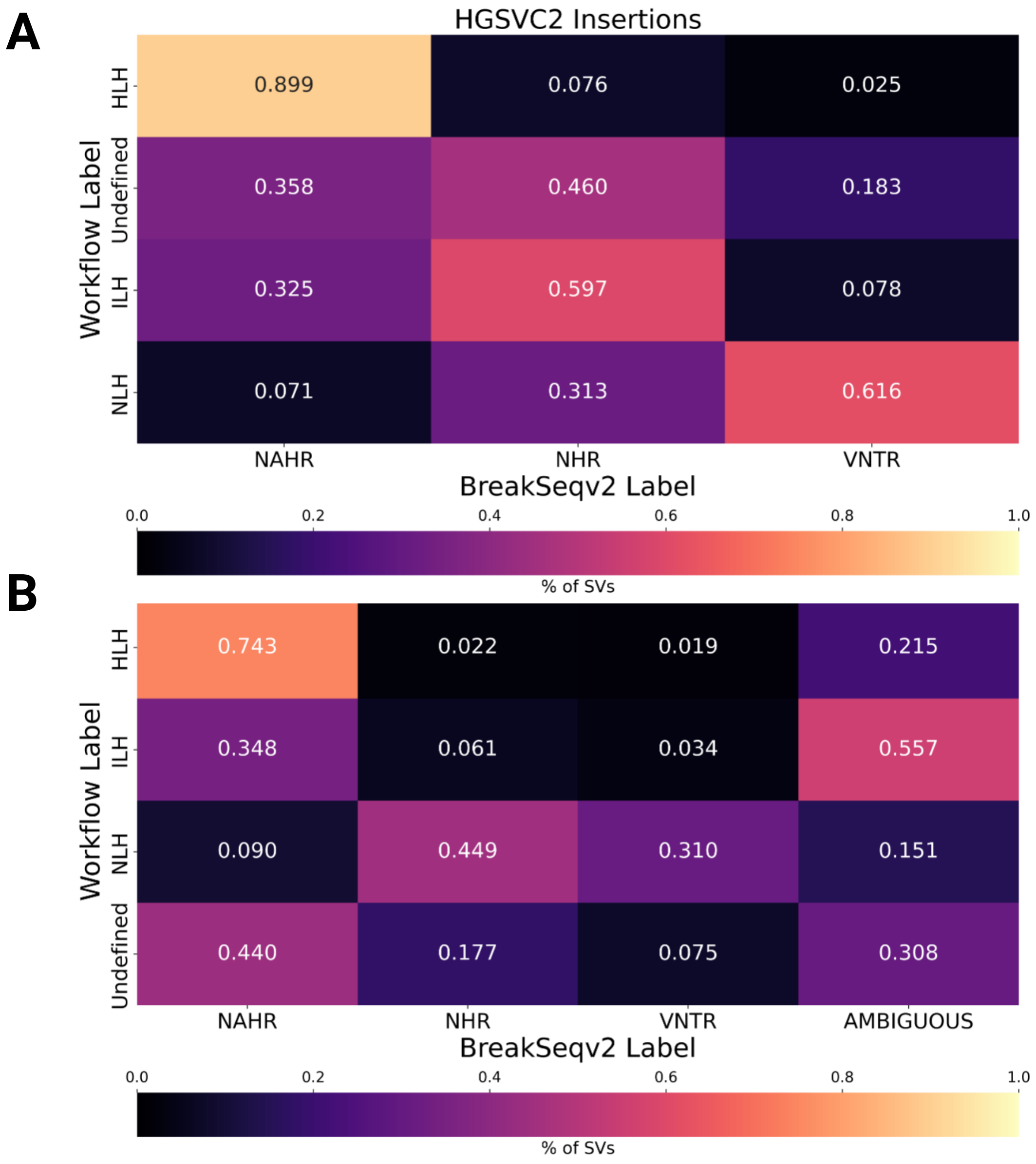


**Figure S3. Mechanism concordance between the homology-based workflow and Breakseq v2.** Heatmaps visualized labels applied to HSGCV2 insertions **(A)** and deletions **(B)**. Percent of concordance was calculated row-wise.

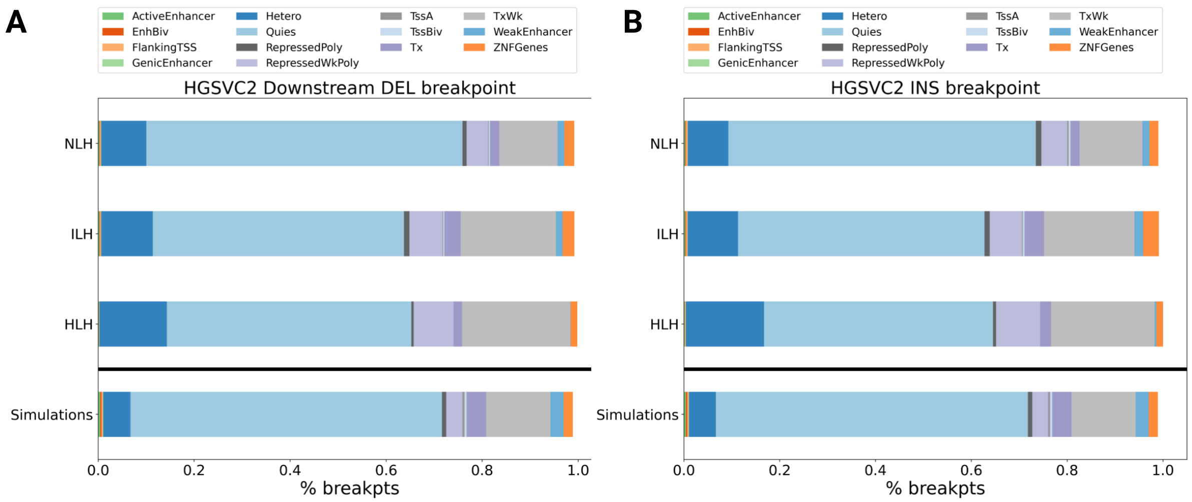


**Figure S4. Intersection of HGSVC2 breakpoints with ChromHMM states**. Stacked barplots depict the proportion of downstream deletion breakpoints **(A)** and insertion breakpoints **(B)** HGSVC2 falling within ChromHMM states. The bottommost bar represents these proportions across the simulations. The bars above show the label specific (HLH, ILH, NLH) breakdown.

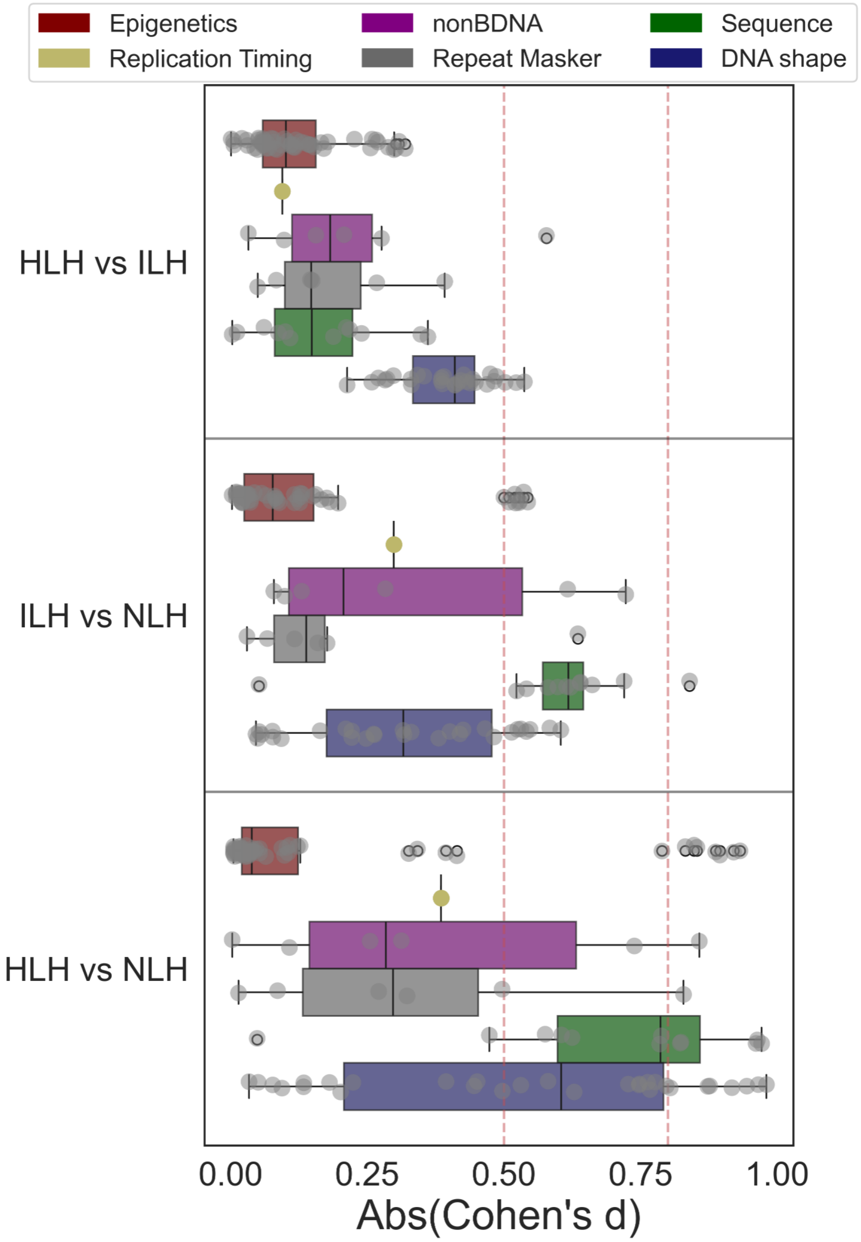


**Figure S5. Underlying features which discriminate homology-based labels, among HGSVC2 insertions.** Boxplots depicting insertions' absolute effect sizes. Each boxplot describes a pairwise comparison between two workflow-labels for a feature type. Effect sizes measured with Cohen's d. Medium and large effect sizes are shown with red dotted lines at 0.5 and 0.8, respectively.

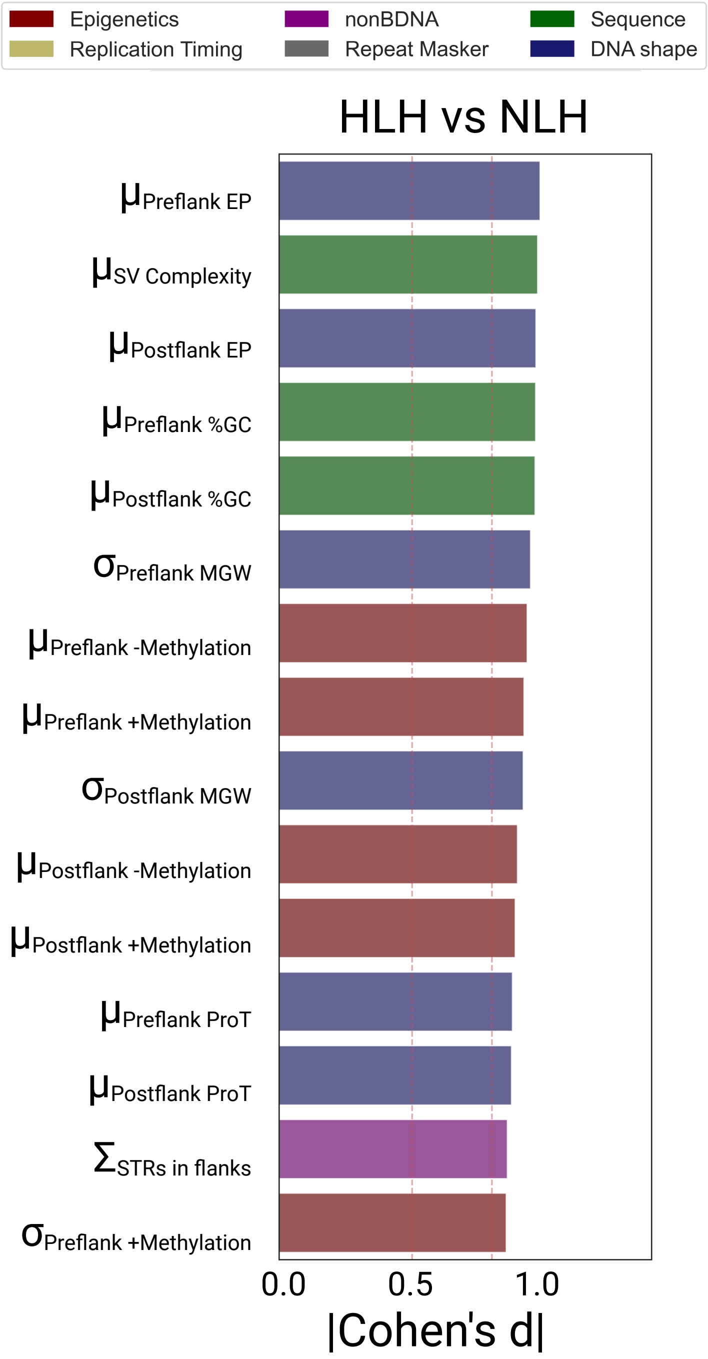


**Figure S6. Underlying features which discriminate homology-based labels, among HGSVC2 insertions.** Barplot highlighting the top 15 most discriminatory features, as per the effect sizes between HLH versus NLH. Color of bar corresponds to feature type.

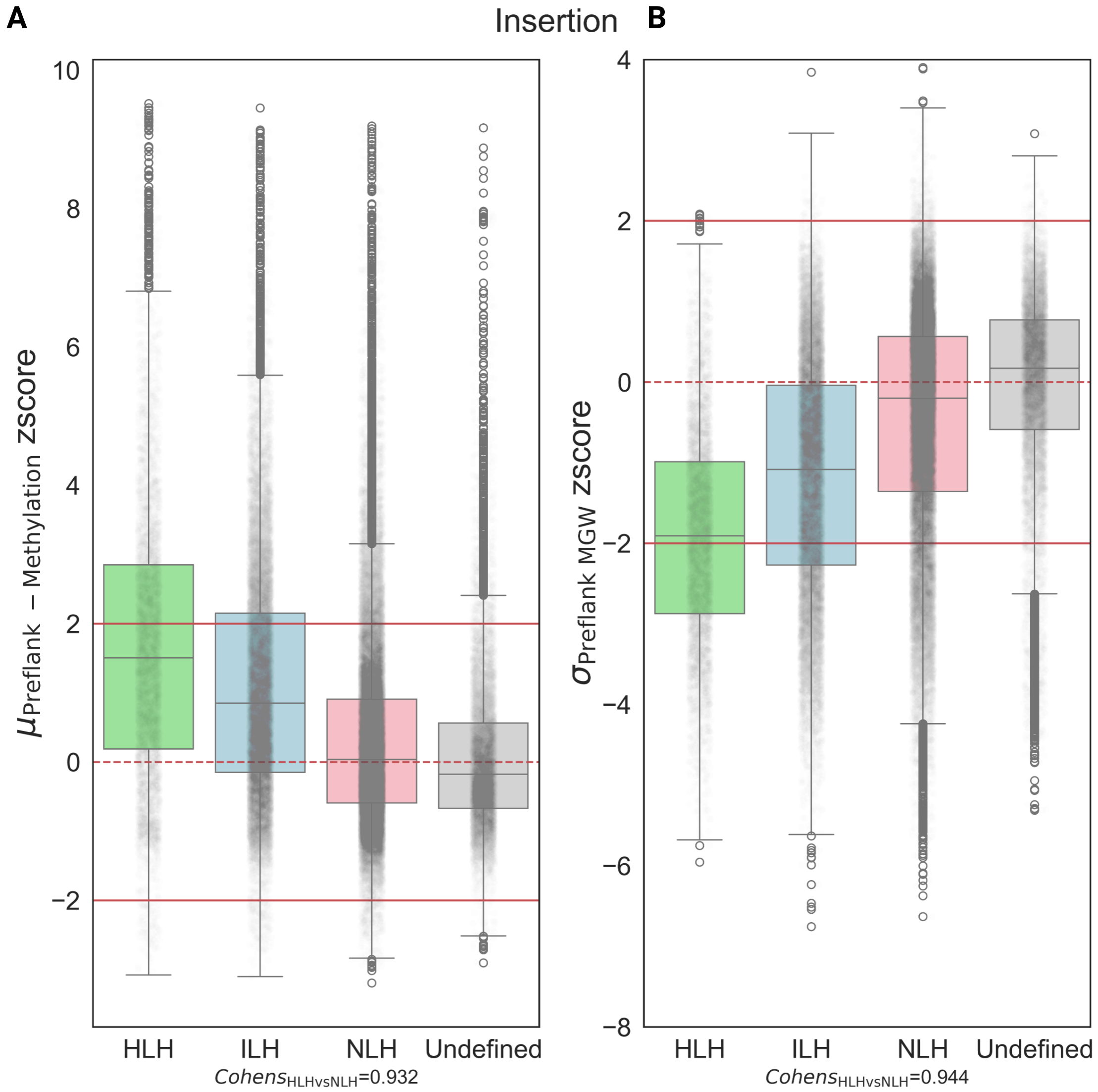


**Figure S7. Underlying features which discriminate homology-based labels, among HGSVC2 insertions.** Boxplots illustrating z-scores by workflow label, as per the mean level of upstream methylation **(A)** and the variation in upstream minor groove width (MGW; **B**).

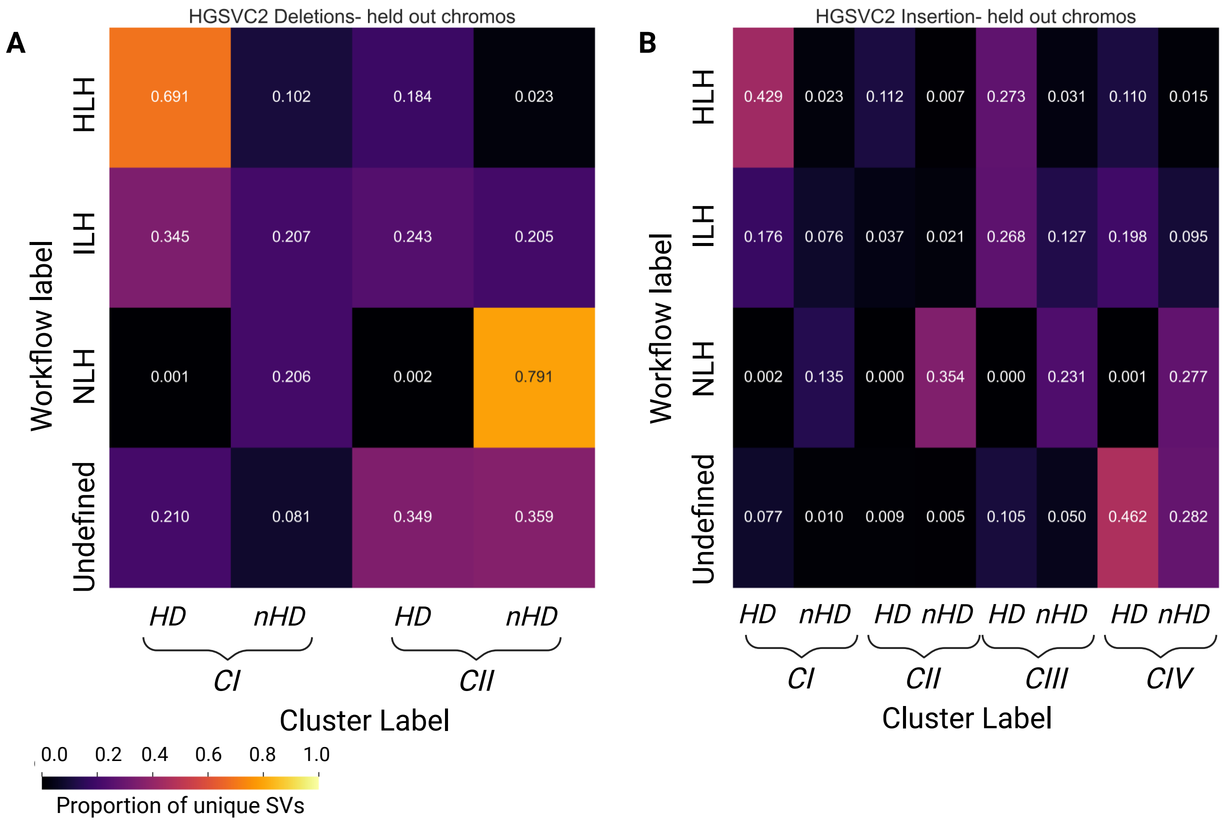


**Figure S8. Performance of cluster-based strategy to held-out datasets.** Heatmap depicting the concordance between the homology-based labels and subcluster-based labels. Heatmaps depict the performance amongst held-out chromosomes from HGSVC2 deletions **(A)** and insertions **(B).**

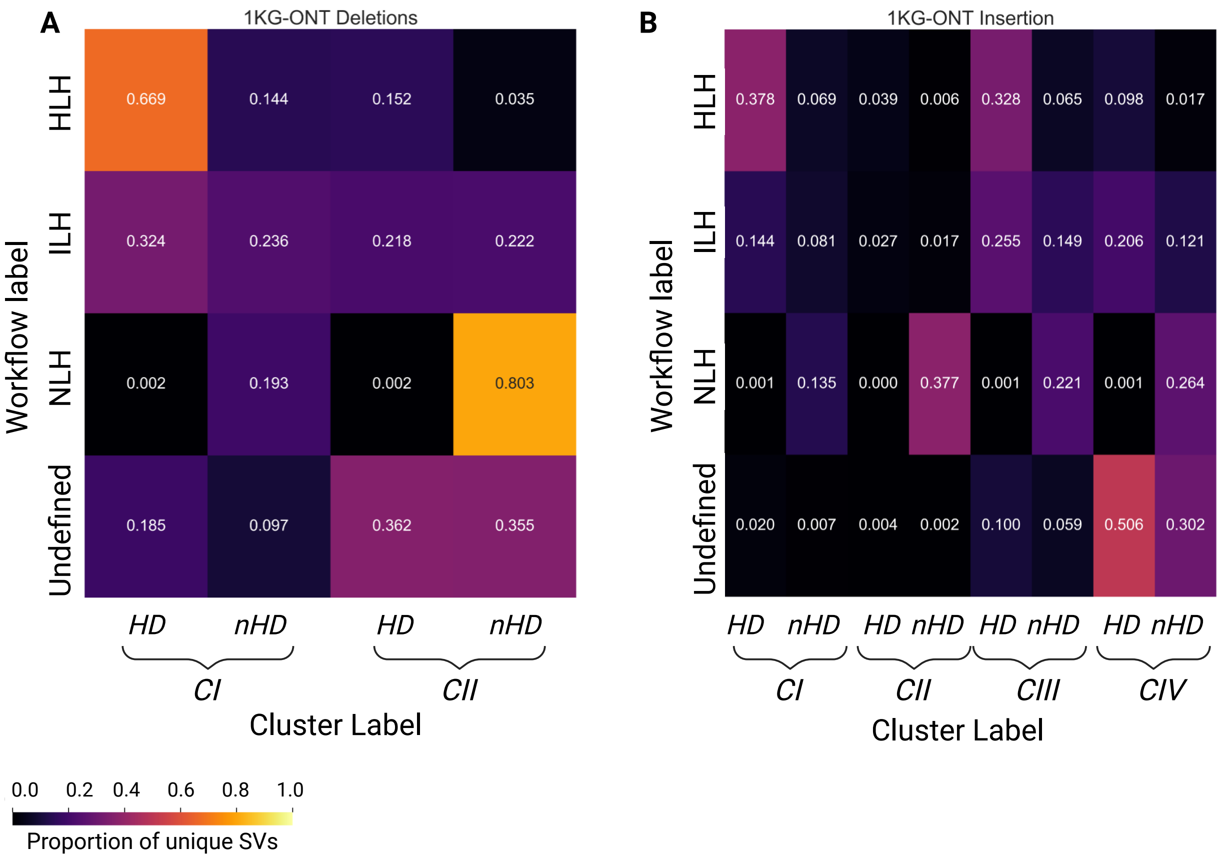


**Figure S9. Performance of cluster-based strategy to held-out datasets.** Heatmap depicting the concordance between the homology-based labels and subcluster-based labels. Heatmaps depict the performance amongst 1KG-ONT deletions **(A)** and insertions **(B).**

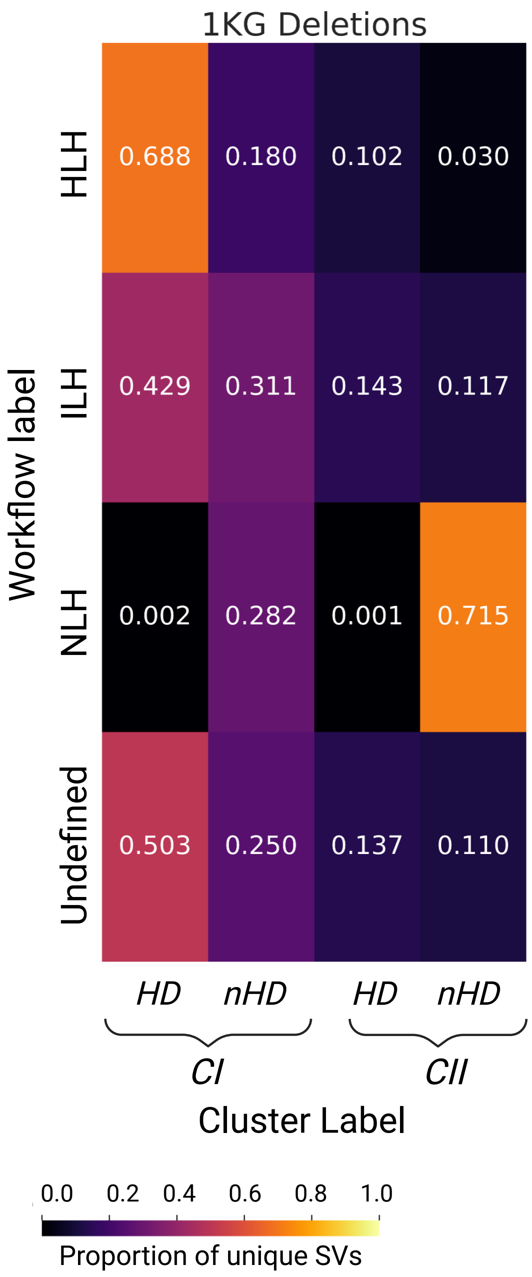


**Figure S10. Performance of cluster-based strategy to held-out datasets.** Heatmap depicting the concordance between the homology-based labels and subcluster-based labels. Heatmaps depict the performance amongst 1KG deletions.

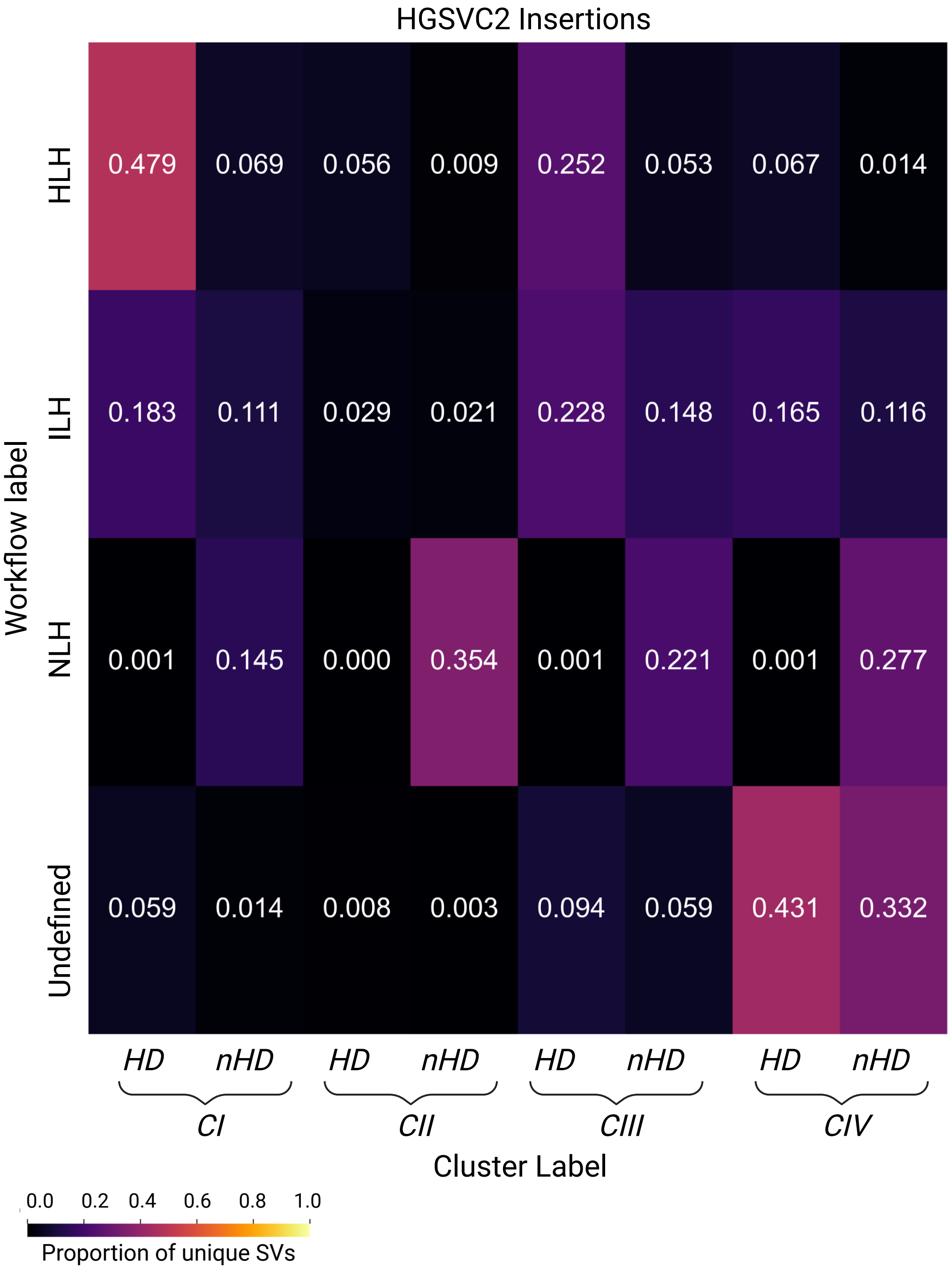


**Figure S11. Performance of cluster-based strategy.** Heatmap depicting the concordance between the homology-based labels and subcluster-based labels. Heatmaps depict the performance amongst HGSVC2 insertions.

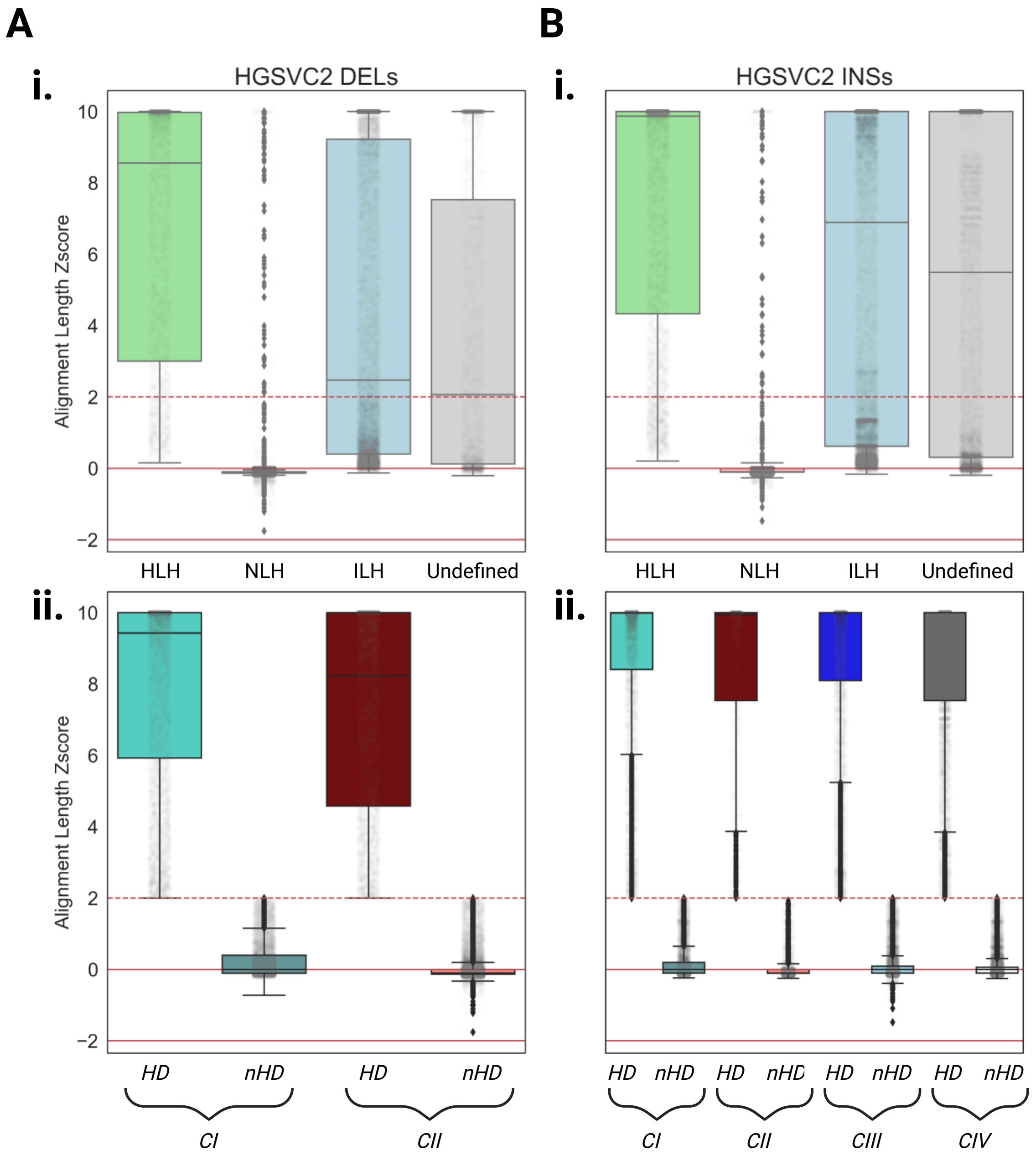


**Figure S12. Preservation of homology patterns following subclustering, among HGSVC2 insertions and deletions.** Boxplots illustrating alignment length z-scores with respect to workflow-based labels for deletions and insertions **(Ai & Bi)**, in comparison to boxplots depicting subclusters **(Aii & Bii)**.

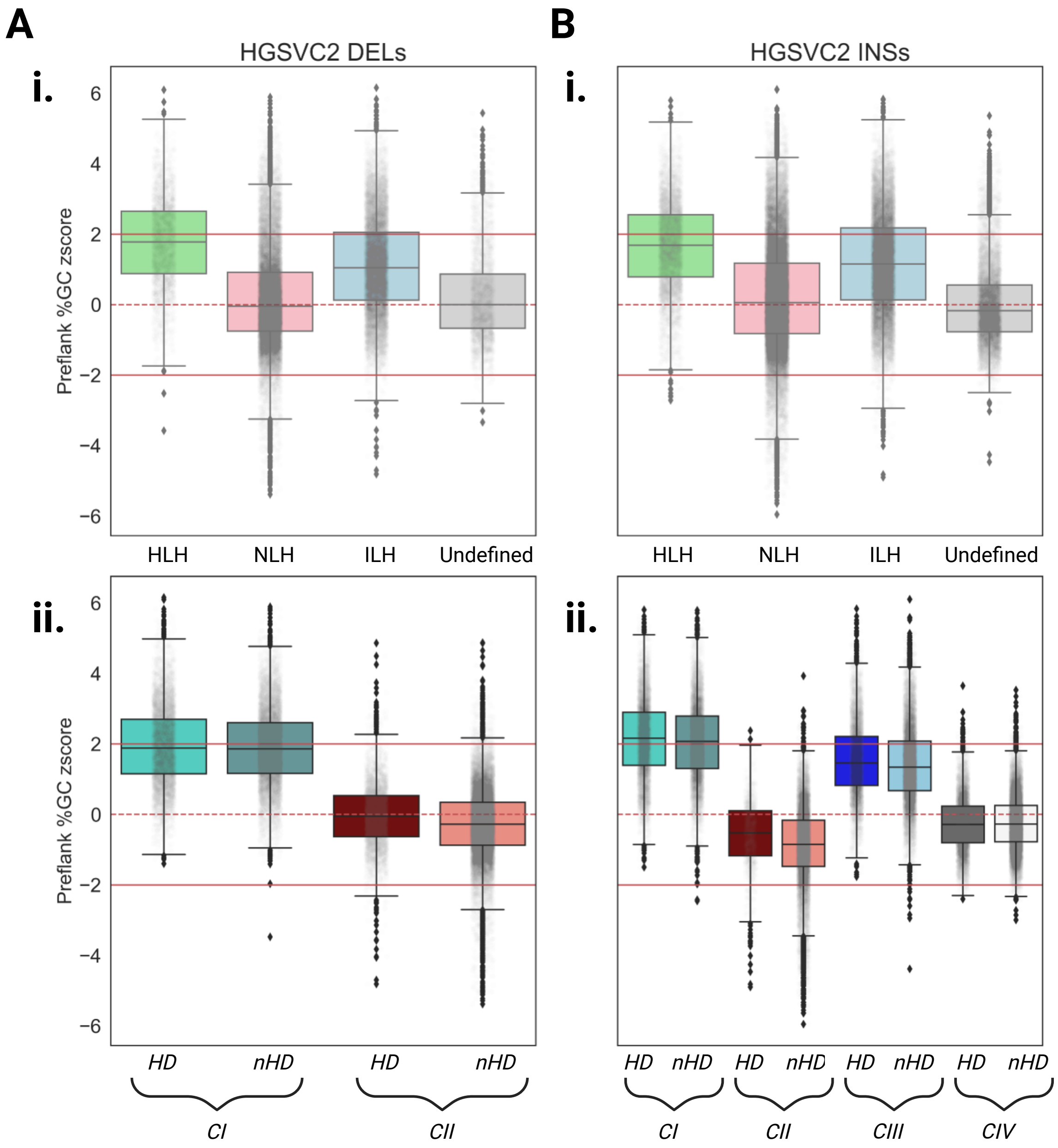


**Figure S13. Preservation of underlying feature patterns following subclustering, among HGSVC2 insertions and deletions.** Boxplots illustrating upstream flanking %GC z-scores with respect to workflow-based labels for deletions and insertions **(Ai & Bi)**, in comparison to boxplots depicting subclusters **(Aii & Bii)**.

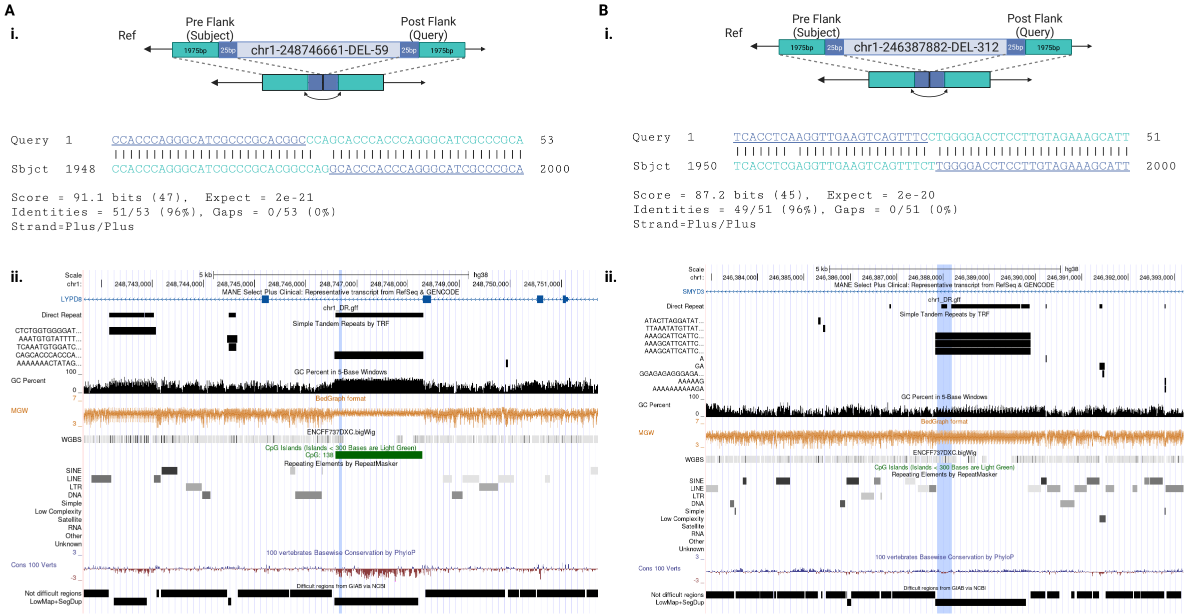


**Figure S14. Underlying sequence features differentiate ILH SVs, subject to clustering**. The raw Blast alignment results from two ILH SVs, chr1-248746661-DEL-59 (**Ai**) and chr1-246387882-DEL-312 (**Bi**), including the breakpoint positions, e-bit score and alignment identities. However, subject to subclustering, chr1-248746661-DEL-59 and chr1-246387882-DEL-312 were assigned to $CI_{HD}$ and $CII_{HD}$, respectively. UCSC Genome Browser tracks depict the 10,000bp local window to the SVs, repetitive elements, %GC content, MGW profile, methylation from H1-hESCs, genomic conservation, and problematic regions, for chr1-248746661-DEL-59 (**Aii**) and chr1-246387882-DEL-312 (**Bii**).

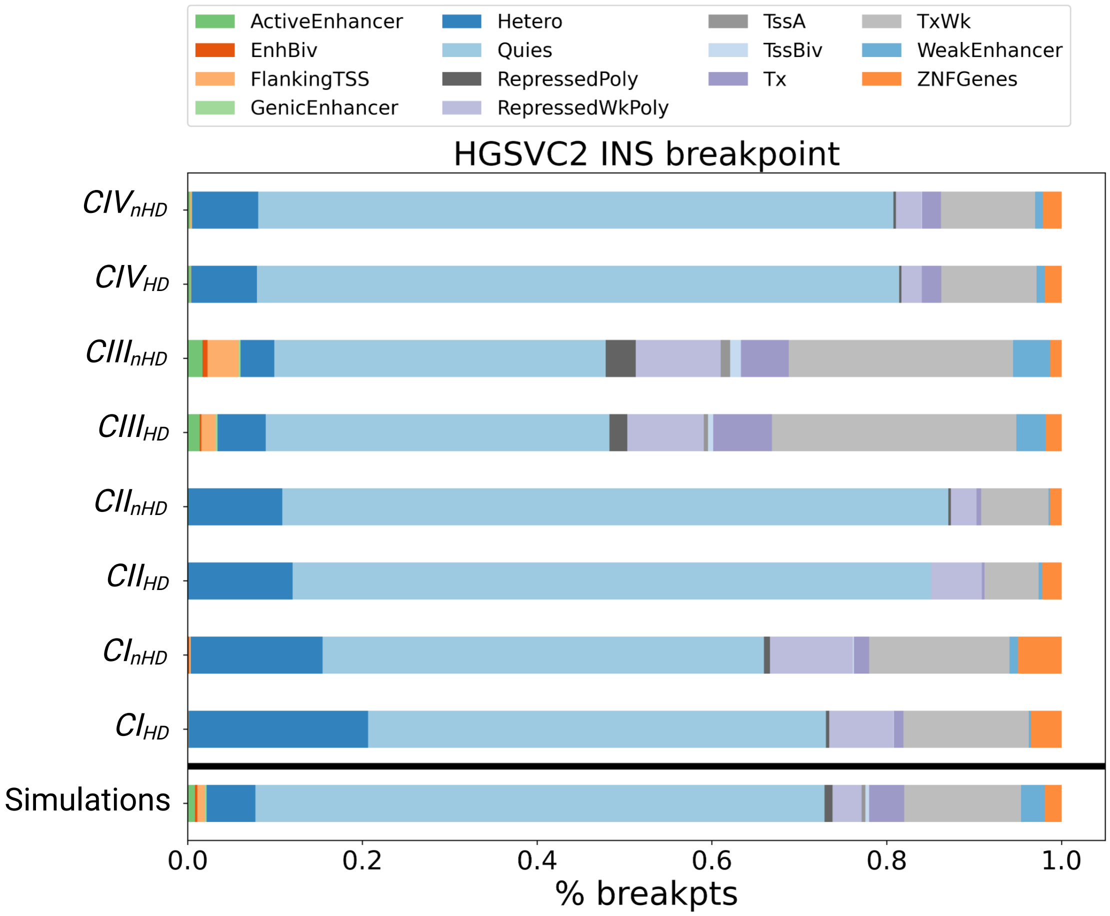


**Figure S15. Intersection of HGSVC2 breakpoints with ChromHMM states**. Stacked barplots depict the proportion of insertions' breakpoints from HGSVC2 falling within ChromHMM states. The bottommost bar represents these proportions across the simulations. The bars above show the cluster specific (clusters *I* to *IV*, along with *HD* and *nHD* subclusters) breakdown.

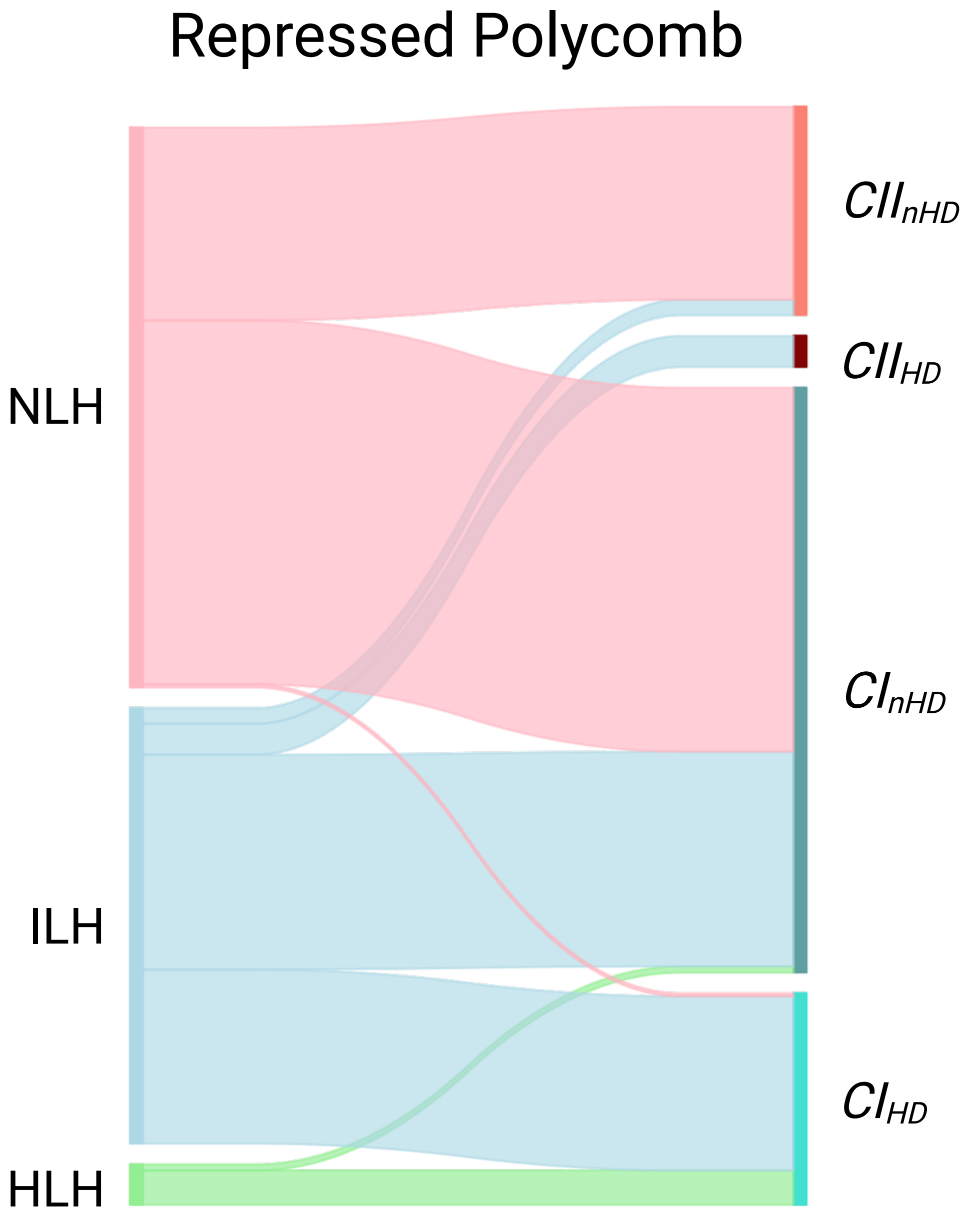


**Figure S16. Sankey plot describing insertions at ChromHMM's repressed polycomb complex from upstream deletion breakpoints.** The left shows the homology-based label, while the right corresponds to the subcluster-based label.

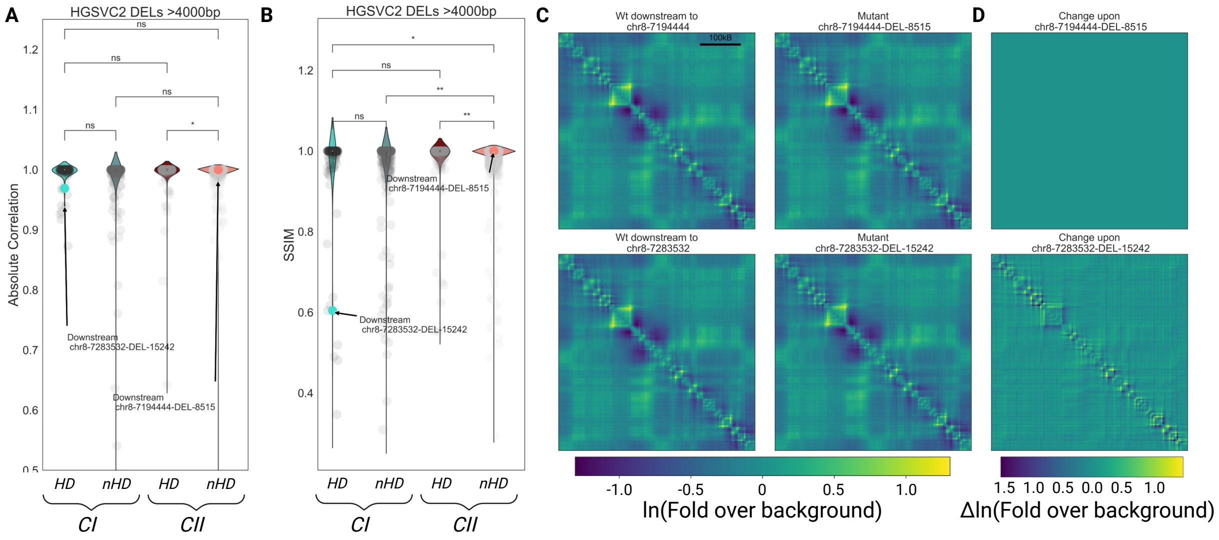


**Figure S17. Cluster1HD SVs appear to alter local 3D genomic folding. (A-B).** Boxplots comparing subclusters' SVs' global and local changes to genomic folding, as per the absolute Pearson's correlation coefficient **(A)**, and structural similarity index measure (SSIM; **B**). (Mann-U Whitney). The $CI_{HD}$ SV, chr8-7283532-DEL-15242, and the $CII_{nHD}$ SV, chr8-7194444-DEL-8515, are annotated on the plots with arrows and coloured markers. **(C-D)**. Heatmap visualizations of the change to folding using these two SVs, whereby the $CII_{nHD}$ SV is depicted along the top row, and the $CI_{HD}$ SV is on the bottom row. The wildtype and mutant 500Mb downstream contact maps are in the first and second column respectively **(C)**. Finally, the change in local DNA interactions upon the $CII_{nHD}$ and $CI_{HD}$ SVs within the downstream window are directly compared **(D)**.

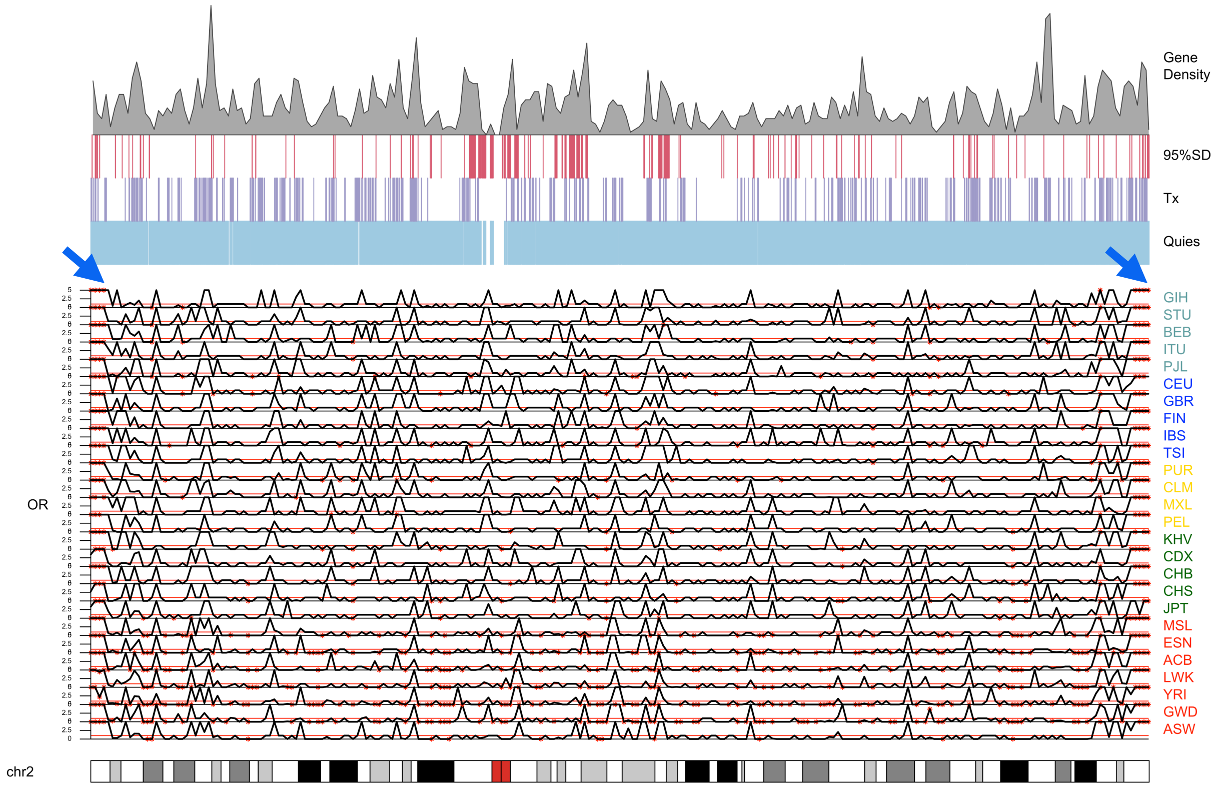


**Figure S18. Consistent $CI_{HD}$ hotspots are found in the subtelomeres, regardless of the chromosome, depicted with chromosome 2 hotspots.** Karyoplots of SV hotspots across 1Mb windows among rare deletions in 1KG. The chromosome's ideogram is found along the bottom. Above this, are the odds ratio depicting the enrichment of $CI_{HD}$, is shown with black lines, for all the populations in 1KG. Colour of text correspond to superpopulation. A thin red line depicts a line at 1, demonstrating no enrichment in $CI_{HD}$ or $CII_{nHD}$ in the window. Further, windows with statistical significance, after p-value adjustment, are depicted with red stars. A blue arrow points to windows with consistent and significant hotspots across ancestries. Above these plots, are genomic elements aligned to the chromosome. These elements include gene density, 95% segmental duplications (SD), along with strong transcription (Tx) and quiescent (Quies) sites from ChromHMM.

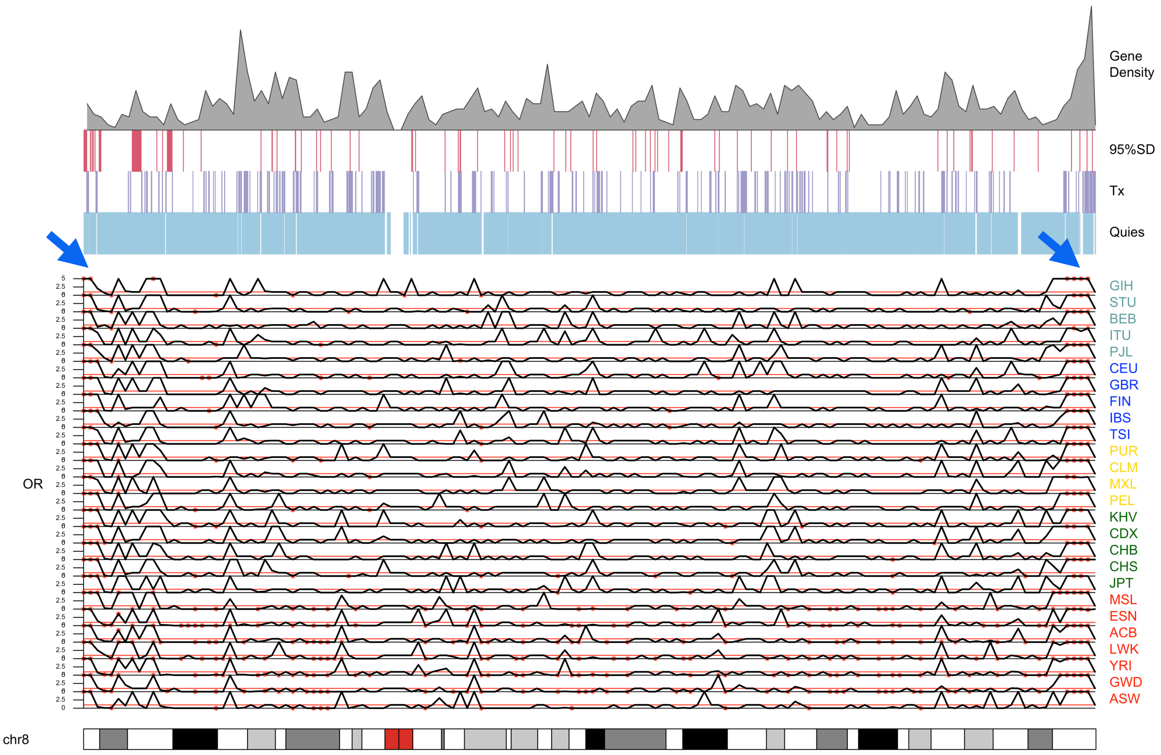


**Figure S19. Consistent $CI_{HD}$ hotspots are found in the subtelomeres, regardless of the chromosome, depicted with chromosome 8 hotspots.** Karyoplots of SV hotspots across 1Mb windows among rare deletions in 1KG. The chromosome's ideogram is found along the bottom. Above this, are the odds ratio depicting the enrichment of $CI_{HD}$, is shown with black lines, for all the populations in 1KG. Colour of text correspond to superpopulation. A thin red line depicts a line at 1, demonstrating no enrichment in $CI_{HD}$ or $CII_{nHD}$ in the window. Further, windows with statistical significance, after p-value adjustment, are depicted with red stars. A blue arrow points to windows with consistent and significant hotspots across ancestries. Above these plots, are genomic elements aligned to the chromosome. These elements include gene density, 95% segmental duplications (SD), along with strong transcription (Tx) and quiescent (Quies) sites from ChromHMM.
